# Novel hyperplastic expansion of white adipose tissue underlies the metabolically healthy obese phenotype of male LFABP null mice

**DOI:** 10.1101/2025.04.07.647661

**Authors:** Anastasia Diolintzi, YinXiu Zhou, Angelina Fomina, Seema Husain, Labros S. Sidossis, Susan K. Fried, Judith Storch

## Abstract

Obesity is an important risk factor for the development of metabolic syndrome disorders. We previously showed that the liver fatty acid-binding protein null mouse (LFABP^−/−^) becomes obese upon high-fat diet (HFD) feeding but remains metabolically healthy. Here we find that the obese LFABP^−/−^ mouse increases subcutaneous adipose tissue (SAT) mass by markedly increasing the number rather than the size of adipocytes, as is typical with HFD. Indeed, while HFD-fed LFABP^−/−^ mice had almost double the fat mass of WT, SAT adipocyte size was >4-fold smaller and adipocyte number was 5-fold higher in the LFABP^−/−^. Transcriptomic analysis of SAT revealed that *Lfabp* deletion alters the expression of multiple pathways that modulate adipose expansion and function including cholesterol biosynthesis, adipogenesis, and extracellular matrix remodeling. LFABP is expressed in liver and small intestine but not in adipose tissues, thus its ablation may promote interorgan crosstalk that drives hyperplastic expansion of metabolically beneficial SAT, contributing to the healthy obese phenotype of the LFABP^−/−^ mouse.

## 1. Introduction

Liver fatty acid-binding protein (LFABP; FABP1) is a highly abundant intracellular protein in liver and small intestine [1]. In addition to long-chain fatty acids (LCFAs), LFABP binds other lipid species including lysophospholipids, monoacylglycerols (MAGs), fatty acyl-CoAs, endocannabinoids (eCBs), bile acids (BAs), and prostaglandins [2]. When challenged with a high-fat diet (HFD), LFABP null (LFABP^−/−^) mice become substantially more obese compared to their wild-type (WT) counterparts, but they maintain a metabolically healthy obese (MHO) phenotype characterized by normoglycemia, normoinsulinemia, decreased hepatic steatosis and increased spontaneous physical activity [3]. Intriguingly, HF-fed LFABP^−/−^ mice, despite their greater adiposity, are also protected from a HFD-induced decline in exercise capacity, displaying an approximate doubling of running distance and time to exhaustion compared with WT mice [4]. Associated metabolic alterations include elevated plasma free fatty acid (FFA) levels post-exercise [4], which may be indicative of increased adipose tissue lipolysis to fuel the exercising muscle.

During overweight/obesity development, adipose tissue (AT) expands through a combination of adipocyte hypertrophy, *i.e.*, an increase in size of existing cells, and hyperplasia, *i.e.,* recruitment, proliferation, and differentiation of new adipocyte progenitor cells (APCs) in a process called adipogenesis [5–7]. Recruitment of inflammatory cells and vascular and extracellular matrix (ECM) remodeling occur as well, allowing sufficient tissue expansion, oxygen supply and nutrient mobilization [8–11]. Hyperplastic AT growth is considered to be the preferred mechanism of expansion since it protects against metabolic disease by maintaining normal adipocyte size and function, with sufficient lipid storage capacity within AT [12, 13]. An inability to recruit new adipocytes leads to enlargement of the preexisting adipocytes, which is thought to be associated with macrophage infiltration, inflamed and dysfunctional AT, ectopic lipid deposition in non-adipose tissues, as well as local and systemic insulin resistance (IR), all contributing to disease development [14]. In general, the mechanisms by which obesity drives hyperplastic instead of hypertrophic adipose tissue expansion remain poorly understood.

Excess energy storage is normally accomplished by subcutaneous white adipose tissue (sWAT); when its storage capacity is exceeded, excess calories accumulate in visceral WAT (vWAT) depots and ectopic sites such as the liver and muscle, causing lipotoxic insult [15, 16]. Hence, homeostatic remodeling of AT expansion becomes dysfunctional in the context of hypertrophic obesity and sustained energy surplus, with increased adipocyte turnover, adipocyte IR, excess ECM deposition (*i.e.,* fibrosis), reduced angiogenic remodeling, and infiltration of immune cells, thus shaping a proinflammatory and fibrotic milieu [8, 11, 17].

LFABP null mice show marked increases in subcutaneous and visceral fat mass relative to WT [3], however nothing is known regarding tissue cellularity or the mechanism of expansion of these fat depots, nor is there information about brown adipose tissue (BAT) in the LFABP^−/−^ mouse. In the present study we sought to characterize the effects of *Lfabp* ablation on AT during diet-induced obesity (DIO) development using histological, transcriptomic, and physiological analyses, to better understand the MHO phenotype of LFABP^−/−^ mice. The results demonstrate a highly unusual hyperplastic expansion of sWAT in the HFD-fed LFABP null mouse, suggesting that AT may be an important determinant of their MHO phenotype.

## 2. Materials and Methods

### (1) Experimental Design

Male mice on a C57Bl/6J background, as previously described [18, 19], were separated into four groups: (a) WT fed a low-fat diet (LFD), (b) WT fed HFD, (c) LFABP^−/−^ fed LFD, and (d) LFABP^−/−^ fed HFD. Mice were maintained on a 12-h light/dark cycle and had unrestricted access to rodent chow (Purina Laboratory Rodent Diet 5015). Mice were weaned onto a chow diet. At 8 weeks of age, WT and LFABP^−/−^ mice were housed 2-3 per cage and fed either a 45 kcal % fat diet high in saturated fat (HFD; D10080402, Research Diets, Inc., New Brunswick, NJ), or a 10 kcal % fat diet (LFD; D10080401; Research Diets, Inc., New Brunswick, NJ) for 12 weeks. Diet compositions are detailed in Table S1. The level of 45% fat by calories was chosen as it is commonly used to promote obesity in rodents without lowering carbohydrates to levels that would promote ketogenesis [3]. Body weights were recorded weekly. Food efficiency was calculated by dividing the body weight gained on a weekly basis by the amount of weekly food intake per animal, multiplied by the calories per gram of experimental diet (3.9 kcal/g LFD and 4.7 kcal/g HFD) to give the weekly caloric intake consumed (*i.e.,* Food efficiency = Weekly BW gained/(Weekly food intake * Calories/g diet)]. Fat and fat-free mass measurements were obtained by Magnetic Resonance Imaging (MRI) (Echo Medical Systems, LLC., Houston, TX) 3 days prior to starting the feeding intervention and 3 days prior to euthanasia. All animal experimental procedures were approved by the Rutgers University Animal Care and Use Committee.

### (2) Histology and Adipocyte Morphometric Analysis

Adipose tissues from 16-hour fasted animals were dissected rapidly and fixed in 10% neutral buffered formalin (NBF, Avantik Biogroup). Fixed samples were embedded in paraffin, sliced in 5-µm-thick sections, and stained with hematoxylin and eosin (H&E). For collagen detection, sections were stained with Masson’s trichrome reagent. Immunostaining was performed by Research Pathology Services (Rutgers University Biomedical Research Innovation Cores, Piscataway, NJ). Images were acquired with an Olympus VS120 virtual slide microscope and processed with OlyVIA Ver.2.9.1 viewer software. Morphometric analysis of white and brown adipocytes was performed with Fiji/ImageJ 2.1.0/1.53c software [20] and its plugin Adiposoft 1.16 [21] was additionally used for semi-automatic evaluation of white adipocytes, whereas brown adipocytes were evaluated manually. For each fat depot, morphological data were collected from at least 200-300 adipocytes per animal from non-overlapping random fields. Dead adipocytes surrounded by crown-like structures (CLSs) were not sized. Adipocyte size quantification was performed considering adipocytes to have a spherical shape and using the respective formulas for fat cell volume (FCV) and subsequent calculations. Specifically for white adipocytes, fat cell number was calculated as FCN (millions) = [(depot weight (g) * 0.8)/(mean lipid (ug)/cell], where mean lipid (ug)/cell = (FCV (pL) * 0.915)/1000, and triacylglycerol (TAG) density = 0.915 g/mL [13, 22, 23].

### (3) Preparation of tissue and RNA isolation

At the end of the feeding period, mice were food deprived for indicated times, anesthetized with ketamine/xylazine/acepromazine (80:100:150 mg/kg, intraperitoneally, respectively), and tissues collected. Mice were euthanized under anesthesia. Inguinal, and epididymal white and interscapular brown adipose tissue (iWAT, eWAT, and iBAT, respectively) depots from mice fasted for 4h were treated with RNA*later*^TM^ stabilization solution (Invitrogen^TM^, Thermo Fisher Scientific, Waltham, MA) upon excision before snap freezing in liquid nitrogen (N_2_) and storage at −80°C.

Total mRNA was extracted from iWAT and iBAT depots using the PureLink^TM^ RNA Mini Kit (Invitrogen^TM^, Thermo Fisher Scientific, Waltham, MA). All tissue samples were first homogenized using TRIzol^TM^ reagent (Invitrogen^TM^, Thermo Fisher Scientific, Waltham, MA) for total RNA isolation. RNA abundance and quality were assessed using a NanoDrop 2000 Spectrophotometer (Thermo Fisher Scientific, Waltham, MA). Two micrograms of total RNA were reverse transcribed using the High-Capacity cDNA Reverse Transcription Kit with RNase Inhibitor (Applied Biosystems^TM^, Thermo Fisher Scientific, Waltham, MA).

### (4) RNA Sequencing Analysis

Total cellular mRNA was extracted as described above and RNA extracts derived from the iWAT of WT and LFABP^−/−^ mice were submitted for RNA-Sequencing (RNA-Seq) analysis to the Genomics Center of Rutgers New Jersey Medical School (Newark, NJ). RNA quality was first checked for integrity using an Agilent 2200 TapeStation (Agilent Technologies); samples with RNA integrity number (RIN) >7.0 were used for subsequent processing. Total RNA was subjected to two rounds of poly(A) selection using oligo-d(T)25 magnetic beads (New England Biolabs). Illumina compatible RNA-seq library was prepared using a NEB next ultra RNA-seq library preparation kit. The cDNA libraries were purified using AmpureXP beads and quantified on an Agilent TapeStation and Qubit 4 Fluorometer (Thermo Fisher Scientific). An equimolar amount of barcoded libraries was pooled and sequenced on the Illumina NovaSeq platform (Illumina, San Diego, CA) using the 1X100 cycles configuration. CLC Genomics Workbench 20.0.4 version (http://www.clcbio.com/products/clc-genomics-workbench/; Qiagen) was used for RNA-seq analysis. De-multiplexed fastq files from RNA-Seq libraries were imported into the CLC software. Bases with low quality were trimmed and reads were mapped to the reference Mus musculus genome GRCm38. The reference genome sequence and annotation files were downloaded from ENSEMBLE, release.92 (Mus_musculus.GRCm38.92.fa, and Mus_musculus.GRCm38.92.gtf). The aligned reads were obtained using the RNA-Seq Analysis Tool of CLC Genomics Workbench using default settings (GEO Series number: **GSE277001**). Comparison of samples was performed using the Gene Set Enrichment Analysis (GSEA) software (v20.3.x), provided from the Broad Institute (https://www.gsea-msigdb.org/gsea/index.jsp), run on the GenePattern platform (https://www.genepattern.org/) and following the GSEA User Guide (http://www.gsea-msigdb.org/gsea/doc/GSEAUserGuideFrame.html) [24, 25]. From the Metabolic Signatures Database (MSigDB, v7.5.1) (https://www.gsea-msigdb.org/gsea/msigdb/index.jsp) available, the KEGG, HALLMARK, and REACTOME gene set collections were used, which are coherently expressed signatures derived by aggregating many metabolic signature database (MSigDB) gene sets to represent well-defined biological states or processes [26, 27]. For each gene set a normalized enrichment score (NES) and a false discovery rate (FDR) q-value were generated. Five independent replicates for each group were used for analysis of differential expression. Differentially expressed pathways and genes with expression values >20 **<u>R</u>**eads **<u>P</u>**er **<u>K</u>**ilobase per **<u>M</u>**illion mapped reads (RPKM), FDRq value <0.1 and fold change |FC| >1.2 were used for downstream analysis.

### (5) Adipose Cholesterol Determination

Adipose samples were homogenized in 10mM phosphate-buffered saline (PBS), as previously described [28]. Lipids were extracted using chloroform-methanol (2:1 v/v) by the method of Folch et al [29]. Lipid extracts from known amounts of tissue and a 5-point concentration gradient using authentic standards were spotted on K5 Silica Gel 150A TLC plates (Whatman #4852-820) and developed using a nonpolar solvent system (hexane-diethyl ether-acetic acid, 70:30:1 v/v). The plates were dried thoroughly and exposed to iodine vapors for 20 min to visualize and identify the lipid spots. Mass densitometry was analyzed using Fiji/ImageJ [20]. Data are expressed as mg cholesterol/g adipose tissue.

### (6) Immunohistochemical (IHC) Staining for Macrophage Infiltration

Sections were deparaffinized, dehydrated through a graded ethanol series, and subjected to heat-induced epitope retrieval with citrate buffer, pH 6.0 for 20 min at 98 °C using a pressure cooker. Primary antibody Rabbit monoclonal anti-F4/80 (Cell Signaling cat# 70076) was applied to sections at dilution of 1:750 for 1 hour followed by an incubation in the secondary antibody Horse anti-Rabbit IgG Polymer (Vector MP6401) for 30 minutes. DAB (3,3’Diaminobenzidine) chromogen substrate (Vector Labs SK-4105) was added for 5 minutes for development of brown color followed by 1 minute in hematoxylin (Vector H-3404) for background blue color.

### (7) Statistical Analysis

The data are presented as mean ± S.D. For body composition, food intake, and adipocyte quality - related measurements, comparisons were made between WT and LFABP null mice for each experimental diet, as well as within the same genotype between LFD and HFD. Differences between the groups were assessed using two-way ANOVA. *Post hoc* comparisons were performed using the Tukey test. For gene expression analysis, comparisons were made only between the HF-fed WT and LFABP null mice using two-tailed unpaired Student’s t-test. The data were analyzed using GraphPad Prism Version 9.5.1 for macOS (GraphPad Software, San Diego, CA, USA). The results were considered statistically significant when *P*<0.05.

## 3. Results

### *Lfabp* deficiency drives hyperplastic expansion of inguinal WAT (iWAT) during obesity development

As shown in Table 1, under LF-feeding LFABP^−/−^ and WT mice have similar body weight (BW), however LFABP^−/−^ mice have 27% higher BW after 12-weeks of HFD (*P* < 0.001) (Fig. 1A). Interestingly, body composition analysis indicated that even on LF-feeding, the percent body fat of LFABP^−/−^ mice was significantly higher compared to their WT counterparts (*P* < 0.001), a difference also found upon HF-feeding (*P* < 0.001) (Fig. 1B). While the percent lean mass is lower when compared with WT mice (*P* < 0.001) (Fig. 1C) absolute lean mass remains unchanged (Table 1). Food efficiency is not different between LF-fed WT and LFABP^−/−^ mice (*P* = 0.919) but is over 40% greater than WT in HF-fed LFABP^−/−^ animals (*P* < 0.001) (Fig. 1D). *Lfabp* ablation resulted in a 65% increase in total BW gain over the 12-week HFD intervention (*P* < 0.001) (Fig. 1E).

**Table 1.** Phenotypic comparison of wild-type and LFABP knockout mice on LFD and HFD.

| Parameter | LFD |  | *P*-Value<br>LFABP<sup>-/-</sup><br>*vs* WT | HFD |  | *P*-Value<br>LFABP<sup>-/-</sup><br>*vs* WT | *p*-value HFD *vs*<br>LFD |  |
| --- | --- | --- | --- | --- | --- | --- | --- | --- |
|  | WT | LFABP<sup>-/-</sup> |  | WT | LFABP<sup>-/-</sup> |  | WT | LFABP<sup>-/-</sup> |
| Week 12 BW, g<br>(*n*=10) | 28.4 ± 1.5 | 30.7 ± 2.3 | $P = 0.212$ | 34.4 ± 2.0 | 43.6 ± 3.9 | $P < 0.001$ | $p < 0.001$ | $p < 0.001$ |
| Weight gain<br>over 12 wk, g<br>(*n*=10) | 4.2 ± 1.7 | 4.8 ± 1.2 | $P = 0.888$ | 10.6 ± 2.6 | 17.5 ± 1.6 | $P < 0.001$ | $p < 0.001$ | $p < 0.001$ |
| Fat Mass, g<br>(*n*=10) | 2.7 ± 1.1 | 6.7 ± 2.2 | $P < 0.001$ | 8.3 ± 1.7 | 15.4 ± 2.3 | $P < 0.001$ | $p < 0.001$ | $p < 0.001$ |
| Fat Mass, % BW<br>(*n*=10) | 10 ± 3.9 | 21.6 ± 5.8 | $P < 0.001$ | 24.2 ± 4.6 | 36.6 ± 3.7 | $P < 0.001$ | $p < 0.001$ | $p < 0.001$ |
| Adiposity<br>Index, % (*n*=10) | 4.8 ± 0.9 | 8.2 ± 1.1 | $P < 0.001$ | 8.2 ± 1.1 | 10.8 ± 0.8 | $P < 0.001$ | $p < 0.001$ | $p < 0.001$ |
| Lean Mass, g<br>(*n*=10) | 23.4 ± 2.1 | 22.9 ± 0.9 | $P = 0.899$ | 25.3 ± 2.1 | 25.8 ± 1.8 | $P = 0.929$ | $p = 0.112$ | $p = 0.005$ |
| Lean Mass, %<br>BW (*n*=10) | 85.1 ± 5.2 | 74.8 ± 5.3 | $P < 0.001$ | 73.7 ± 5.5 | 62.5 ± 4.6 | $P < 0.001$ | $p < 0.001$ | $p < 0.001$ |
| iWAT, g (*n*=10) | 0.4 ± 0.1 | 0.7 ± 0.2 | $P < 0.001$ | 0.8 ± 0.2 | 1.4 ± 0.2 | $P < 0.001$ | $p < 0.001$ | $p < 0.001$ |
| iWAT, %BW<br>(*n*=10) | 1.4 ± 0.3 | 2.4 ± 0.6 | $P = 0.001$ | 2.3 ± 0.7 | 3.2 ± 0.4 | $P < 0.001$ | $p = 0.005$ | $p = 0.003$ |
| iWAT FCV, pL<br>(*n*=5) | 21.4 ± 6.8 | 46.6 ± 16.3 | $P = 0.642$ | 222 ± 64.1 | 45.6 ± 7.8 | $P < 0.001$ | $p < 0.001$ | $p > 0.999$ |
| iWAT FCN,<br>millions (*n*=5-6) | 31.2 ± 15.6 | 20.6 ± 5.4 | $P = 0.244$ | 4.7 ± 2.7 | 23.6 ± 7.8 | $P = 0.001$ | $p = 0.009$ | $p = 0.925$ |
| eWAT, g (*n*=10) | 0.6 ± 0.2 | 0.8 ± 0.3 | $P = 0.070$ | 1.3 ± 0.2 | 1.9 ± 0.2 | $P < 0.001$ | $p < 0.001$ | $p < 0.001$ |
| eWAT, %BW<br>(*n*=10) | 2.1 ± 0.5 | 2.7 ± 0.8 | $P = 0.160$ | 3.7 ± 0.5 | 4.5 ± 0.5 | $P = 0.024$ | $p < 0.001$ | $p < 0.001$ |
| eWAT FCV, pL<br>(n=5) | 52.5 ± 9.7 | 113 ± 27.1 | P = 0.327 | 296 ± 50.4 | 247 ± 92.6 | P = 0.510 | p < 0.001 | p = 0.007 |
| eWAT FCN, millions (n=5) | 16 ± 5.4 | 8.1 ± 1.9 | P = 0.005 | 4.2 ± 0.5 | 8.9 ± 2.4 | P = 0.127 | p < 0.001 | p = 0.974 |
| iBAT, g (n=10) | 0.1 ± 0.0 | 0.2 ± 0.1 | P < 0.001 | 0.1 ± 0.0 | 0.4 ± 0.1 | P < 0.001 | p = 0.301 | p < 0.001 |
| iBAT, %BW (n=10) | 0.3 ± 0.1 | 0.8 ± 0.2 | P < 0.001 | 0.4 ± 0.1 | 0.8 ± 0.2 | P < 0.001 | p = 0.885 | p = 0.609 |
| Brown adipocyte area, um<sup>2</sup> (n=5) | 374 ± 79.7 | 537 ± 89.1 | P = 0.038 | 449 ± 83.1 | 658 ± 90.2 | P = 0.007 | p = 0.528 | p = 0.157 |
| Liver, g (n=10) | 0.9 ± 0.1 | 1.0 ± 0.1 | P = 0.973 | 1.1 ± 0.1 | 1.6 ± 0.3 | P < 0.001 | p = 0.505 | p < 0.001 |
| Liver, %BW (n=10) | 3.7 ± 0.3 | 3.5 ± 0.4 | P = 0.543 | 3.3 ± 0.1 | 3.9 ± 0.5 | P = 0.002 | p = 0.059 | p = 0.044 |
| Gallbladder, g (n=9-10) | 0.019 ± 0.006 | 0.010 ± 0.003 | P = 0.001 | 0.019 ± 0.005 | 0.013 ± 0.004 | P = 0.009 | p = 0.989 | p = 0.611 |
| Food Efficiency, g gained/kcal consumed (n=10) | 0.005 ± 0.002 | 0.006 ± 0.002 | P = 0.919 | 0.012 ± 0.003 | 0.017 ± 0.001 | P < 0.001 | p < 0.001 | p < 0.001 |
| iWAT Cholesterol, mg/g (n=8-9) | N/A | N/A | N/A | 1.61 ± 0.348 | 2.41 ± 0.555 | P = 0.003 | N/A | N/A |

Data are shown as means ± SD. HFD, high-fat diet; WT, Wild Type; LFABP, Liver Fatty Acid-Binding Protein; BW, body weight; iWAT, inguinal white adipose tissue; FCV, fat cell volume; FCN, fat cell number; eWAT, epididymal white adipose tissue; iBAT, interscapular brown adipose tissue. Adiposity Index was calculated as the ratio of the sum of the excised adipose depots over BW after fasting. Statistical analysis was performed by using two-way ANOVA with Tukey's *post hoc* comparisons.

**Figure 1.**
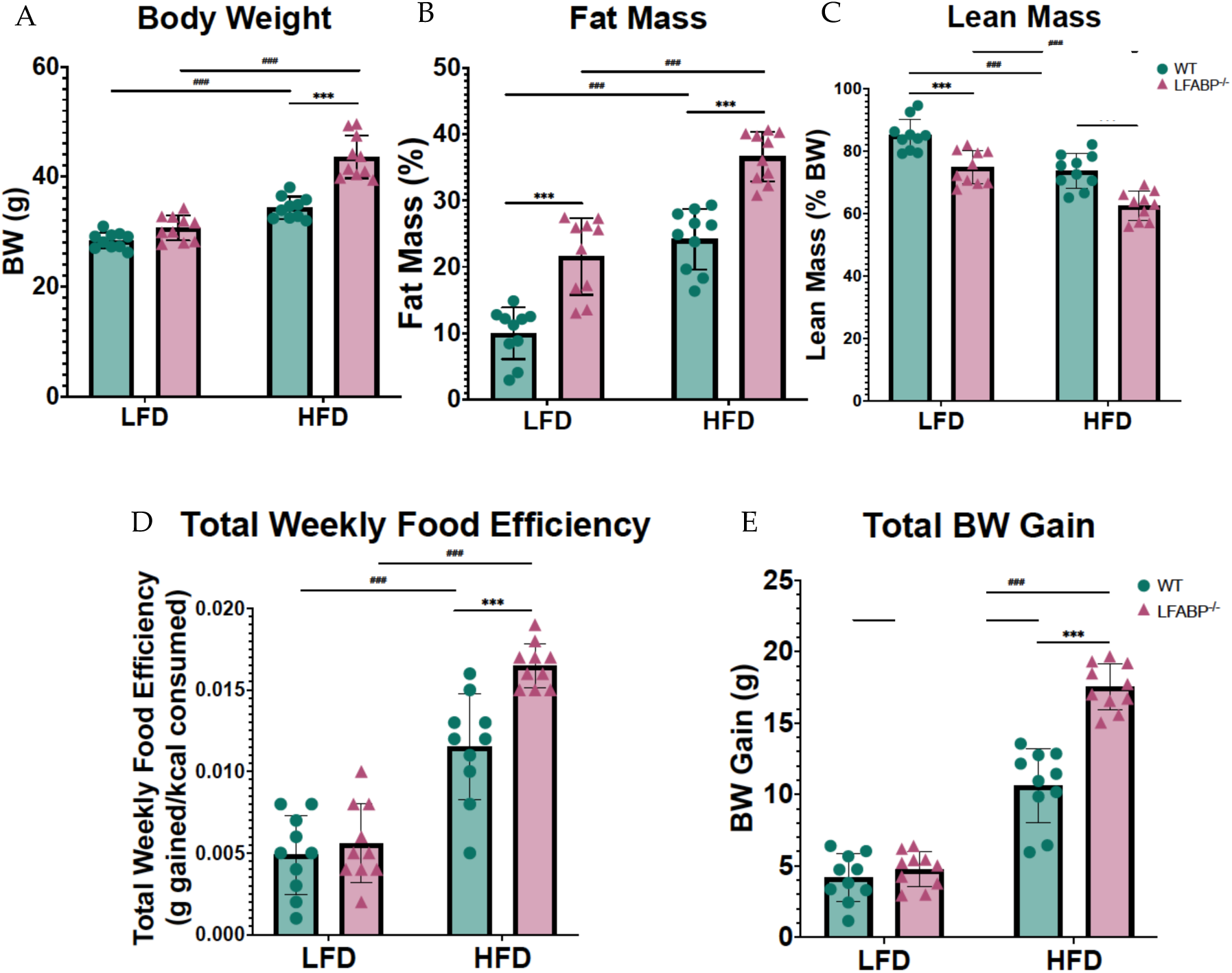
*Lfabp* deletion induces an obese phenotype. **(A)** Post-intervention body weight (g). **(B)** Lean mass (% of total BW). **(C)** Fat mass (% of total body weight, BW). **(D)** Mean weekly food efficiency per mouse (g). **(E)** Total body weight gain (g). n = 10. LFABP*^−/−^ vs.* WT: *\*\*\**, *p* < .0001. HFD *vs.* LFD: *^###^*, *p* < .0001. 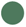 WT; 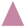 LFABP*^−/−^*. LFD, low-fat diet; HFD, high-fat diet; BW, body weight; WT, wild-type; LFABP*^−/−^*, liver fatty acid-binding protein null.

In this study, iWAT served as a bonafide sWAT and eWAT as a vWAT depot, while iBAT served as a bonafide brown fat depot. On LF-feeding the inguinal white fat depot was 71% heavier in the null mice (iWAT: *P* = 0.001) whereas the epididymal depots were not different (eWAT: *P* = 0.160). Upon HF-feeding, both iWAT and eWAT depots were significantly higher in the LFABP knockout mice (iWAT: *P* < 0.001; eWAT: *P* = 0.024) (Fig. 2, Table 1). Thus, *Lfabp* ablation results in higher absolute levels of iWAT mass which is exacerbated under HF-feeding, reaching almost double the mass of the WT mice (HFD: *P* < 0.001) (Fig. 3A), in agreement with previous results [3]. The iBAT mass was >2-fold higher in the LFABP null mice relative to WT on both diets (*P* < 0.001) (Fig. 2, Table 1).

**Figure 2.**
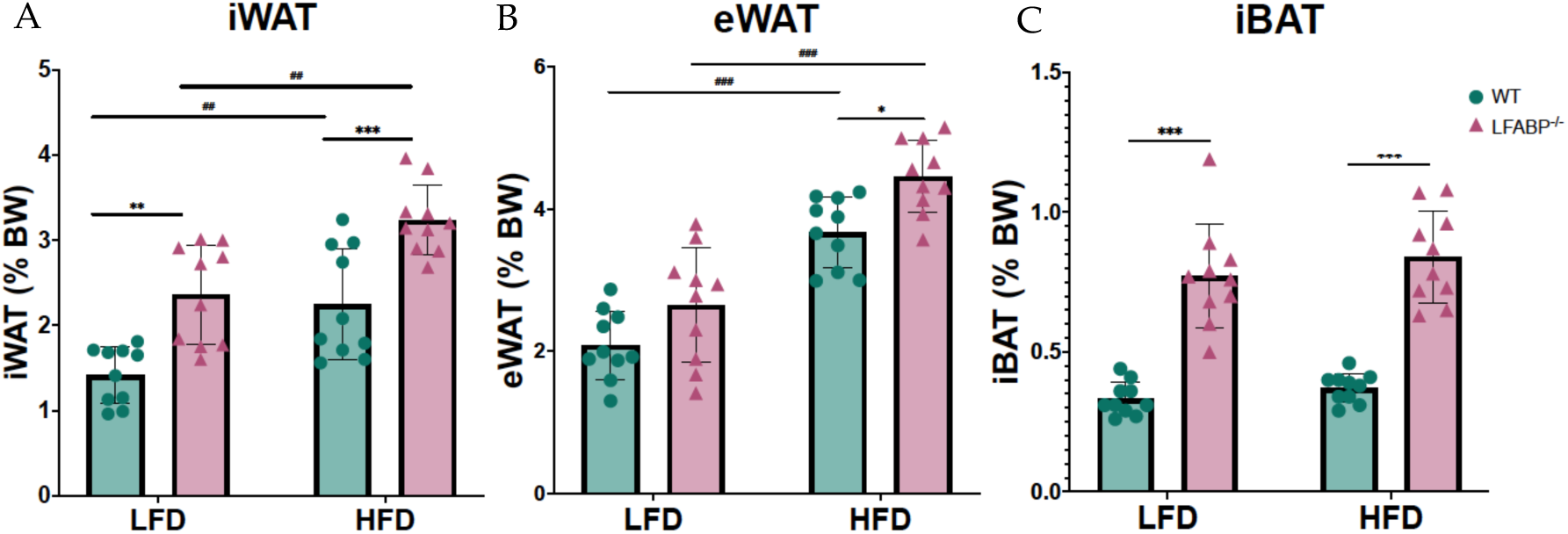
*Lfabp* deletion induces both white and brown adiposity upon DIO. Percentage contribution of **(A)** inguinal white adipose tissue (iWAT), **(B)** epididymal white adipose tissue (eWAT), and **(C)** interscapular brown adipose tissue (iBAT) to body weight (% BW) after LF- and HF-feeding. n = 10. LFABP*^−/−^ vs.* WT: *\*\**, *p* < .001; *\*\*\**, *p* < .0001. HFD *vs.* LFD: *^##^*, *p* < .001; *^###^*, *p* < .0001. 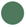 WT; 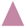 LFABP*^−/−^*. WT, wild-type; LFABP*^−/−^*, liver fatty acid-binding protein null; LFD, low-fat diet; HFD, high-fat diet; BW, body weight; iWAT, inguinal white adipose tissue; eWAT, epididymal white adipose tissue; iBAT, interscapular brown adipose tissue.

**Figure 3.**
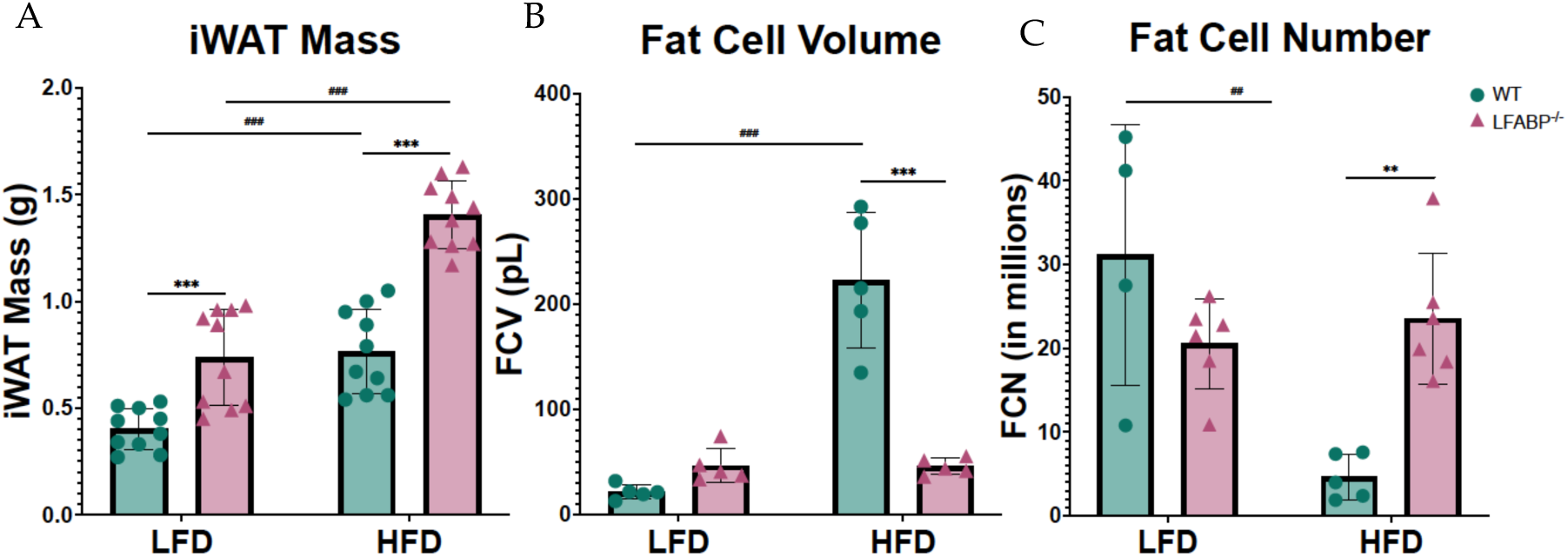
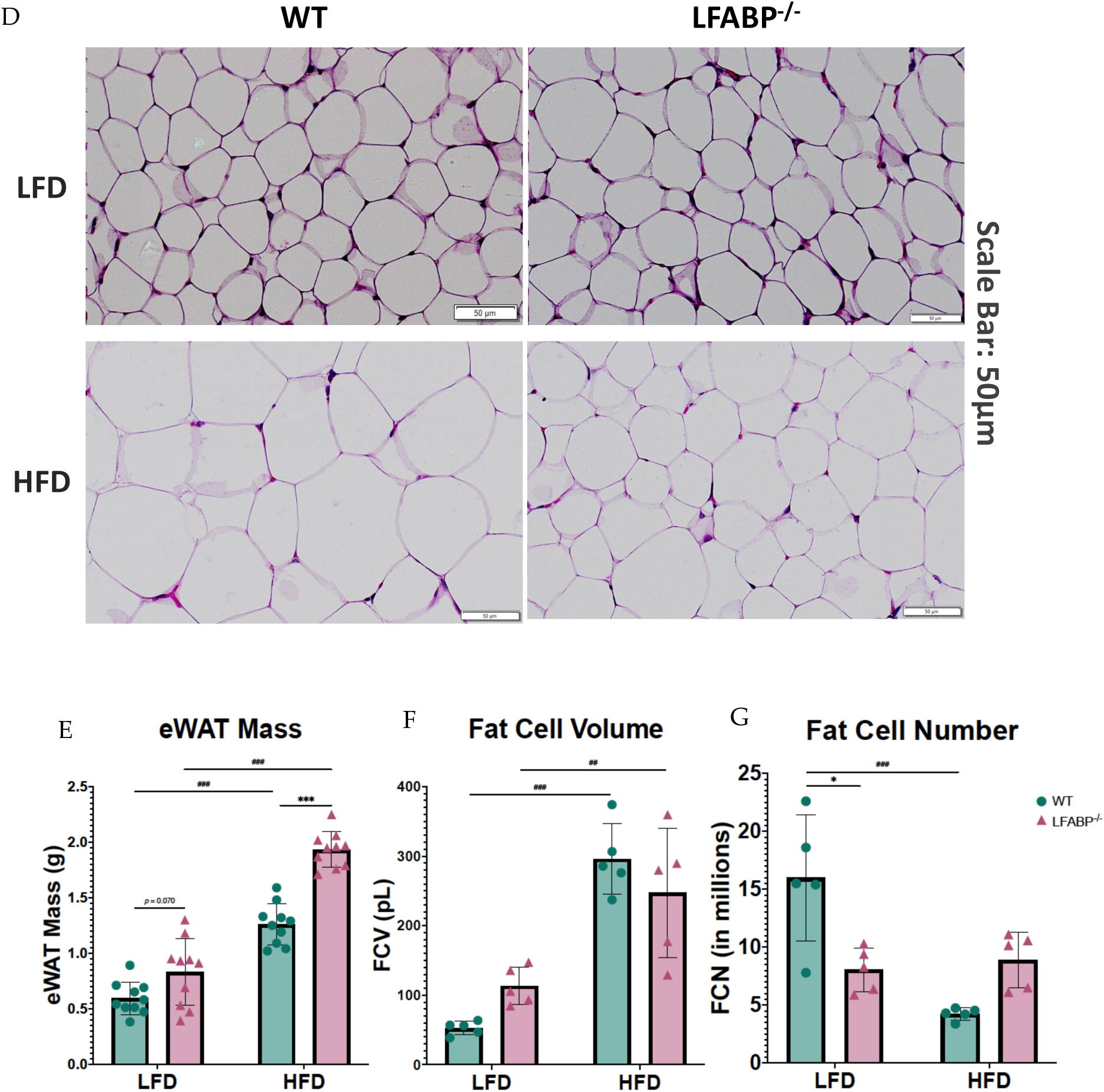
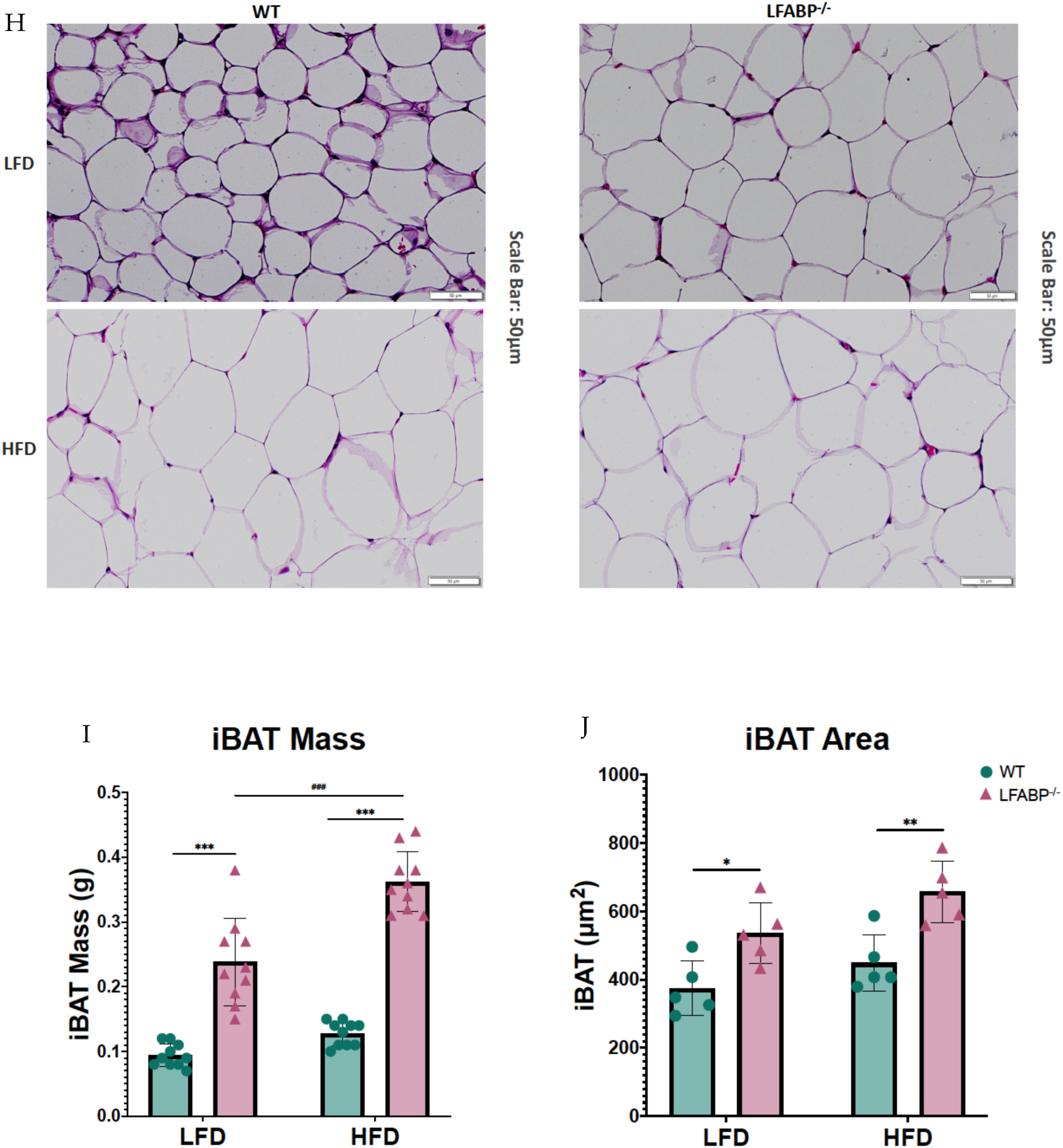
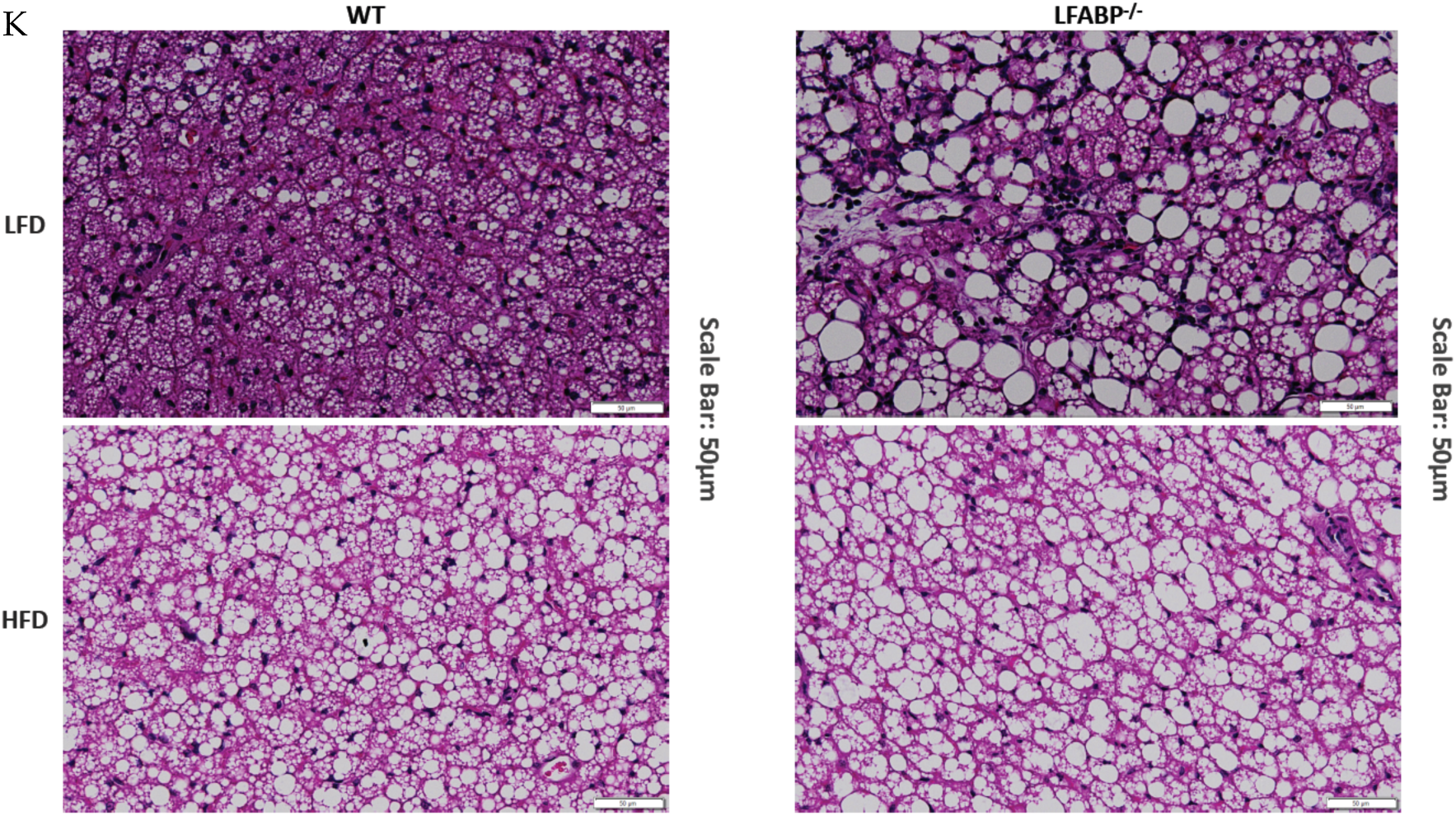
*Lfabp* deletion induces an altered adipocyte phenotype in subcutaneous, visceral, and brown adipose tissue depots. **(A)** Inguinal white adipose tissue (iWAT) mass (g) (n = 10). **(B)** Inguinal fat cell volume (FCV; pL) (n = 5). **(C)** Inguinal fat cell number (FCN) per depot (millions) (n = 5-6). **(D)** Representative H&E-stained pictures of iWAT depots. Scale bar, 50 µm. **(E)** Epididymal white adipose tissue (eWAT) mass (g) (n = 10). **(F)** Epididymal fat cell volume (FCV; pL) (n = 5). **(G)** Epididymal fat cell number (FCN) per depot (millions) (n = 5-6). **(H)** Representative H&E-stained pictures of eWAT depots. **(I)** interscapular brown adipose tissue (iBAT) mass (g) (n=10). **(J)** Brown adipocyte area (µm*^2^*) (n=5). **(K)** Representative H&E-stained pictures of iBAT depots. Scale bar, 50 µm. LFABP*^−/−^ vs.* WT: *\**, *p* < .01; *\*\**, *p* < .001; *\*\*\**, *p* < .0001. HFD *vs.* LFD: *^##^*, *p* < .001; *^###^*, *p* < .0001. 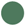 WT; 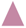 LFABP*^−/−^*. LFD, low-fat diet; HFD, high-fat diet; WT, wild-type; LFABP*^−/−^*, liver fatty acid-binding protein null; iWAT, inguinal white adipose tissue; eWAT, epididymal white adipose tissue; FCV, fat cell volume; FCN, fat cell number; iBAT, interscapular brown adipose tissue.

To determine the cellular basis of the markedly increased adiposity in LFABP^−/−^ mice, we measured cell diameters from each genotype for at least 200 randomly chosen inguinal, epididymal and interscapular brown adipocytes. Diameters for inguinal and epididymal fat cells were converted into cell volumes considering white adipocytes as spheres. LF-fed LFABP^−/−^ and WT mice were found to have comparable inguinal adipocyte size (fat cell volume, FCV; below 50pL/cell) (*P* = 0.642). Upon HF-feeding the increase in iWAT mass is associated with a 4-5-fold increase in adipocyte size in WT mice, as expected (Fig. 3A, B) (*P* <0.001). Surprisingly and in marked contrast to WT, HF-fed LFABP^−/−^ mice maintain an iWAT adipocyte size similar to that on the LFD (*P* > 0.999), and thus have a dramatic 4-fold smaller inguinal adipocyte size (*P* < 0.001) (Fig. 3B) and 5-fold higher numbers of adipocytes in the iWAT depot (*P* = 0.001) (Fig. 3C, D) when compared with their WT counterparts.

LFABP^−/−^ and WT mice have comparable eWAT mass as a % of BW on LF-feeding (*P* = 0.070). While eWAT mass increased significantly for both genotypes when fed a HFD, *Lfabp* ablation results in markedly higher eWAT mass with DIO development (*P* < 0.001) (Fig. 3E). Interestingly, unlike what was found in the iWAT depot, the increase in eWAT mass observed on HF-feeding is associated with an increase in adipocyte size in the LFABP^−/−^ mice, similar to the WT mice (LFABP^−/−^: *P* = 0.007; WT: *P* < 0.001) (Fig. 3F). Notably, however, the increase in eWAT mass on HF-feeding is accompanied by almost double the number of adipocytes in LFABP^−/−^ mice, relative to WT (Fig. 3G, H), although this did not reach statistical significance.

*Lfabp* ablation results in substantially larger iBAT mass and brown adipocyte area, relative to the WT, regardless of feeding intervention (iBAT mass: *P* < 0.001 for both diets; Brown adipocyte area: LFD, *P* = 0.038 and HFD, *P* = 0.007) (Fig. 3I, J, K). Upon HFD feeding, *Lfabp* ablation results in greater iBAT mass relative to LF-feeding (*p* < 0.001), whereas iBAT mass remains unchanged in the WT strain, independent of dietary fat intake (Fig. 3I). It was noted that the brown adipocytes of LFABP null mice appear to be infiltrated with higher lipid content, giving to the whole tissue a lighter appearance (Fig. 3K).

### Differential pathway enrichment in subcutaneous adipose tissue upon *Lfabp* ablation and HFD-induced obesity development

To gain insight into the remarkable histological phenotypes noted in the iWAT, we performed RNA-seq followed by HALLMARK, KEGG, and REACTOME pathway enrichment analyses after 12 wks of HF-feeding. The complete list of up- and down-regulated pathways for all three MSigDBs, as well as the volcano plot for REACTOME pathway enrichment analysis in iWAT of HFD-fed WT and LFABP^−/−^ mice can be found in Supplementary Materials (Tables S2-S3 and Fig. S1).

REACTOME pathway enrichment analysis in iWAT showed that among 1,026 identified pathways (25,254 identified genes), 99 gene sets (137 transcripts) were differentially expressed between LFABP^−/−^ and WT mice (FDRq < 0.1), with 61 upregulated (74 transcripts) and 38 downregulated (63 transcripts) in the HF-fed LFABP^−/−^ mice (|FC|>1.2). We observed consistent upregulation of cholesterol metabolism-related pathways, multiple pathways involved in NF-*κ*B signaling, and mTORC1 and PI3K/Akt/mTOR signaling (Table 2 and Fig. 4A, B, & C). Downregulated gene sets in the HF-fed LFABP^−/−^ iWAT included ECM- and inflammation-related pathways (Table 2 and Fig. 4A, B, & C).

**Table 2.**
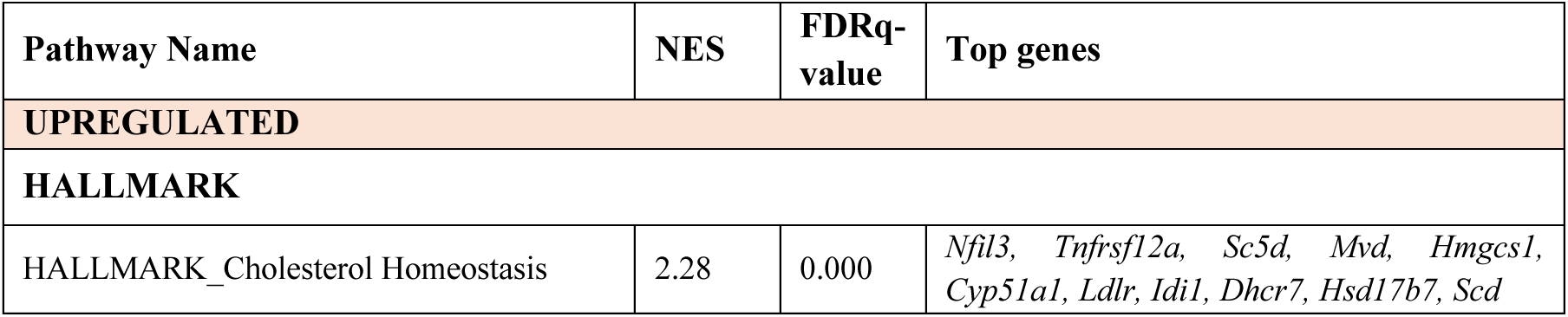
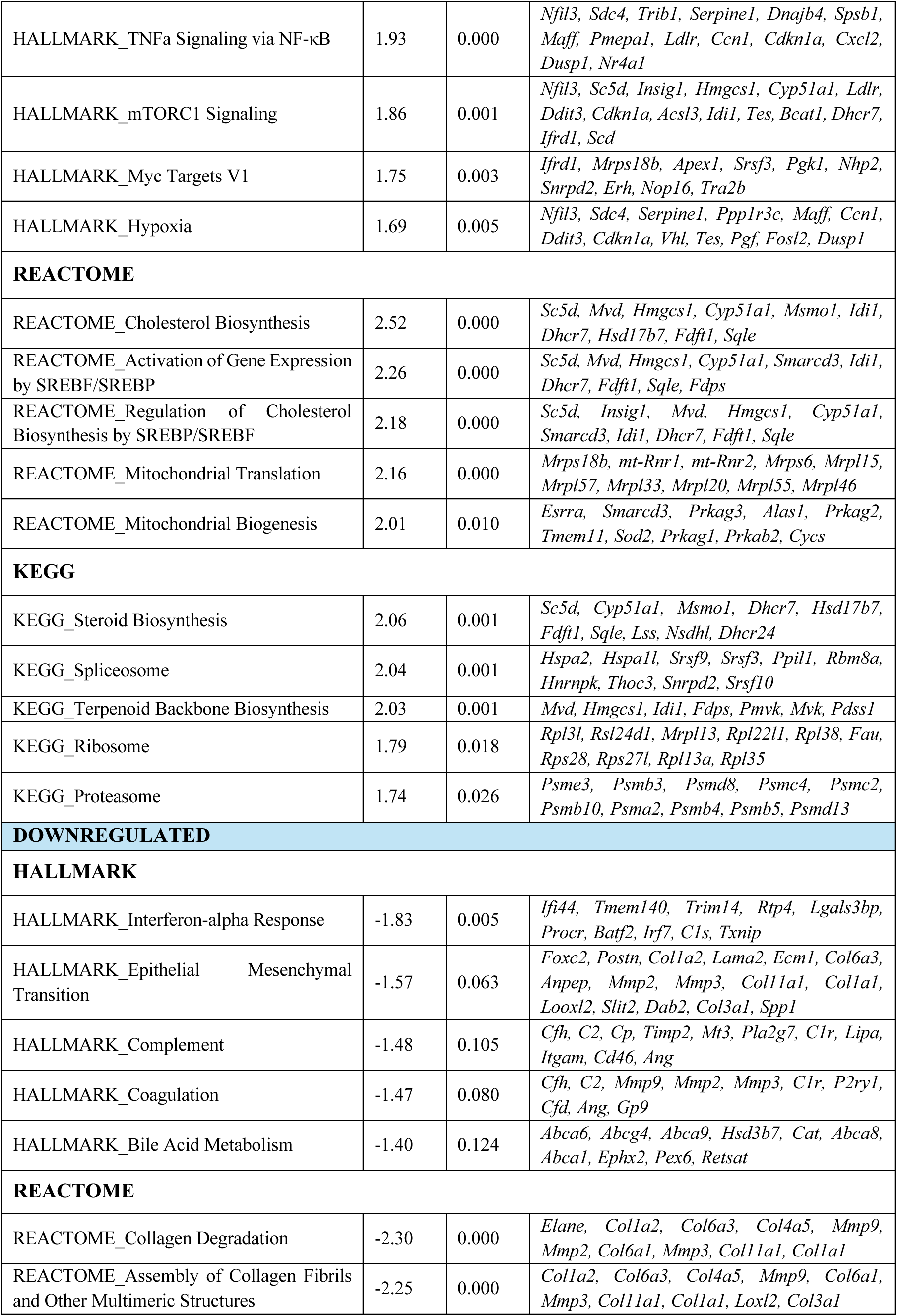
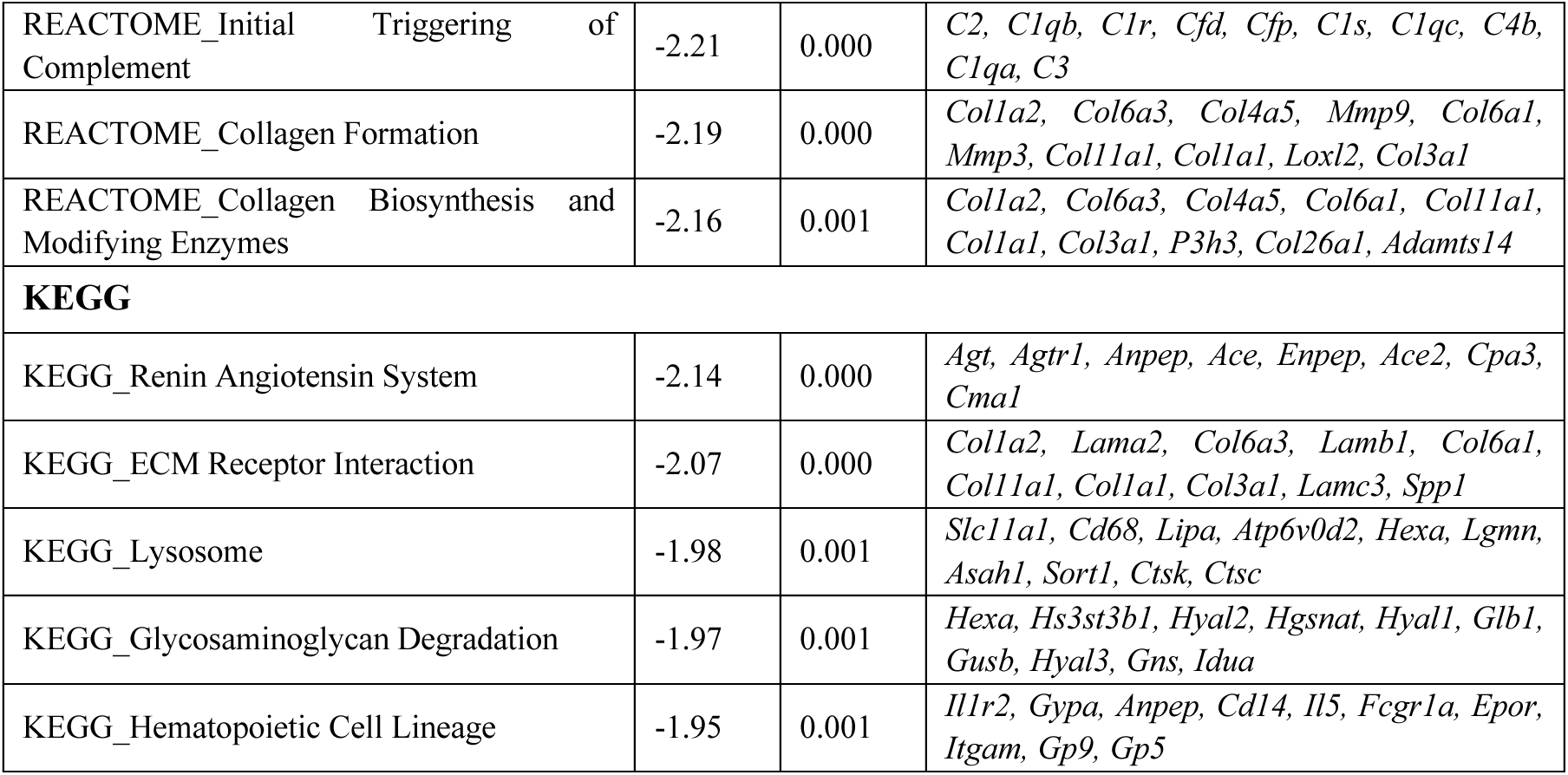
Differential expression of selected gene sets for HALLMARK, REACTOME, and KEGG pathway enrichment analyses in iWAT of LFABP null mice relative to WT.

**Figure 4.**
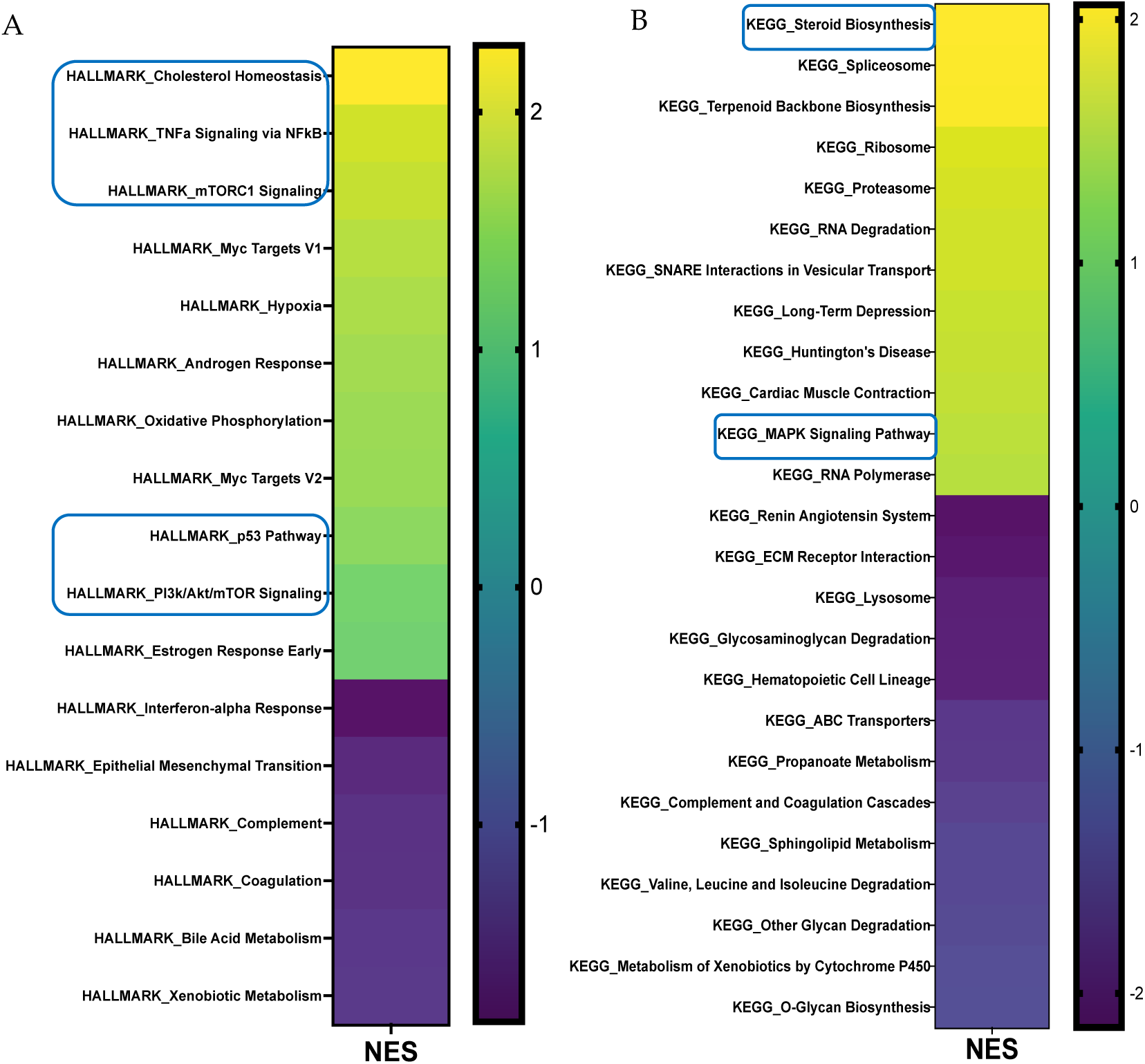
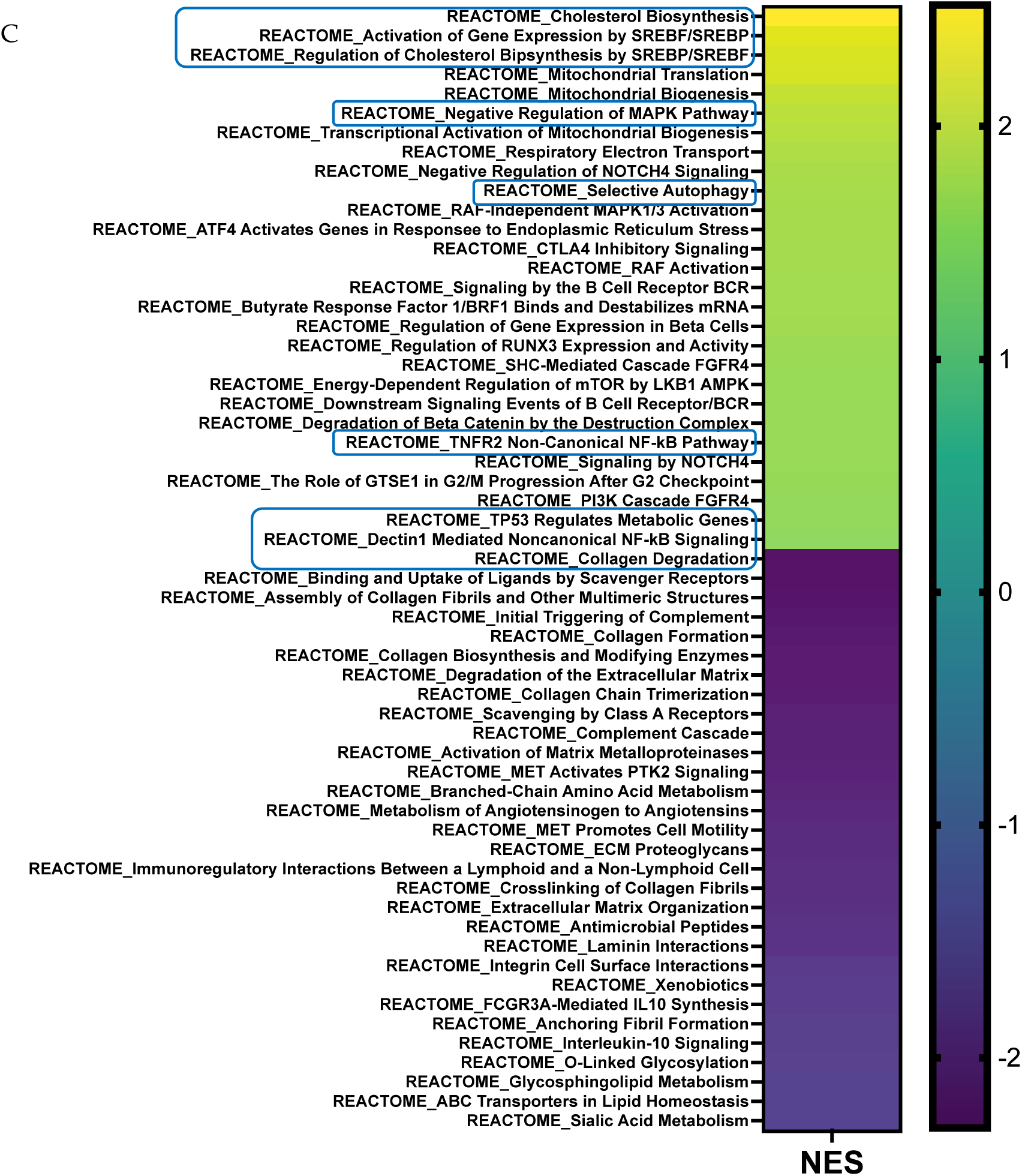
Gene set enrichment analysis (GSEA) of differentially expressed pathways in the iWAT of *LFABP* null mice upon HF-feeding. The heatmaps present pathways that are significantly different for the MSigDB’s **(A)** HALLMARK, **(B)** KEGG, **(C)** and REACTOME (n = 5 / genotype). The direction and significance of each pathway was based on the gene set as a whole, while considering |NES| > 1.2 and FDRq-value < 0.1. The boxes indicate pathways that are common between two or all three MSigDBs. MSigDB, molecular signature database; NES, Normalized Enrichment Score; FDR, False Discovery Rate.

### *Lfabp* ablation induces distinct transcriptional responses in iWAT upon DIO

We sought to evaluate the potential molecular basis for the marked increase in iWAT fat accumulation and unexpected higher cellularity upon obesity development and LFABP deficiency by interrogating the gene expression signature of the iWAT depot to identify changes in genes related to growth and obesity status. Among 25,254 identified genes, 137 transcripts were expressed differentially between LFABP^−/−^ and WT mice (FDRq < 0.1), with a total of 74 upregulated and 63 downregulated in the HF-fed LFABP^−/−^ mice (|FC|>1.2). The top upregulated and downregulated differentially expressed genes are shown in Fig. 5 and Table 2.

**Figure 5.**
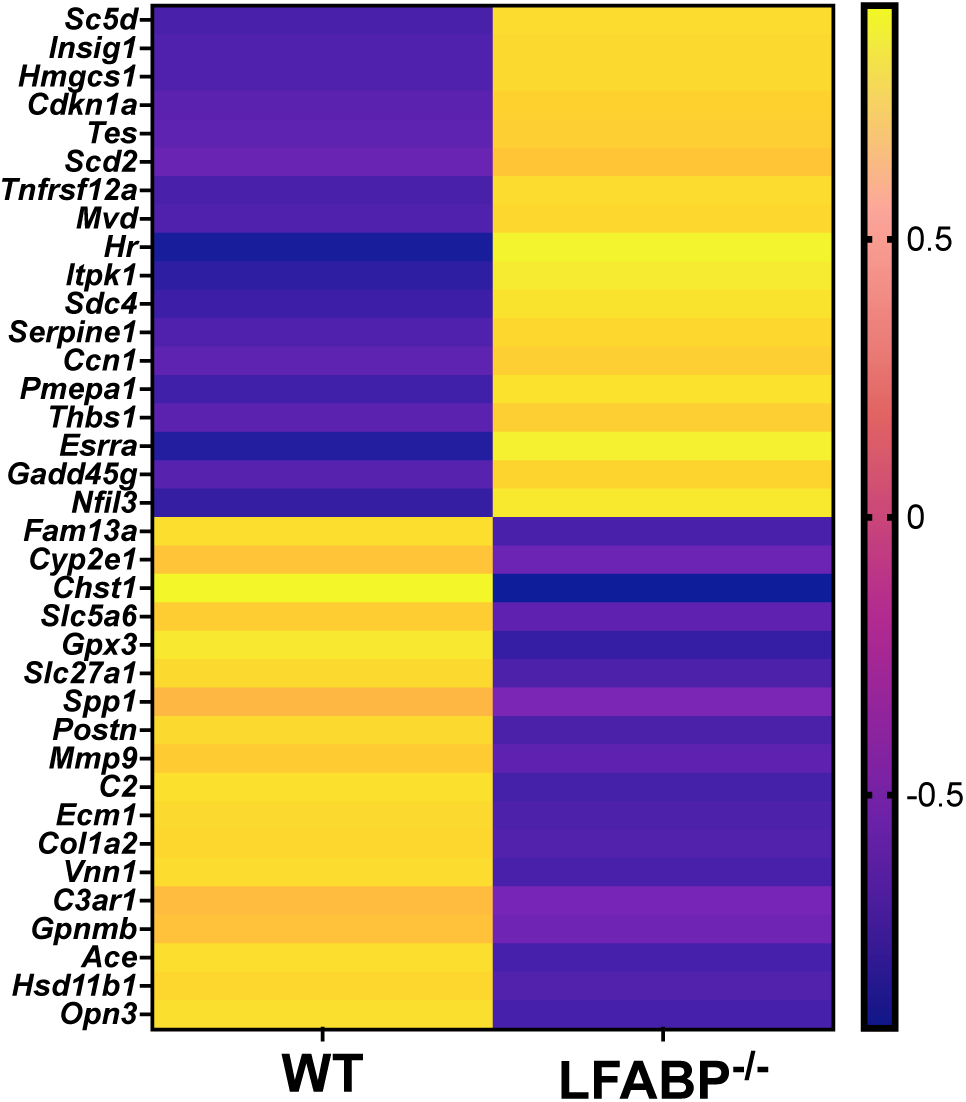
*Lfabp* ablation induces alterations in the transcriptomic profile of iWAT upon HF-feeding. Heatmap of differentially expressed transcripts with RPKM values > 20, FDRq-value < 0.1, and |FC| > 1.2 between LFABP null and WT mice (n = 5 / genotype). FDR, False Discovery Rate; RPKM, Reads Per Million reads mapped.

### *Lfabp* null mice show altered adipogenic potential

Given the striking hyperplastic phenotype observed in the iWAT of the LFABP^−/−^ mouse model, we inspected individual transcripts proposed to denote enhanced adipogenesis. Interestingly, we found approximately 3.5-fold downregulation of cytochrome P450 family 2 subfamily E member 1 (CYP2E1), characteristic of adipogenesis regulator (Areg) cells that inhibit adipogenesis through the retinoic acid signaling pathway [30], as well as those of family with sequence similarity 13 member A (FAM13A), implicated in adipocyte size and fat distribution [31], in the iWAT of HF-fed LFABP^−/−^ mice relative to WT. Additionally, Hairless (*Hr*) expression, which is significantly higher in the LFABP^−/−^ mice upon DIO, is required for white adipocyte development both *in vitro* and *in vivo* and is thought to be proadipogenic [32]. The orphan nuclear receptor estrogen-related receptor α (ESRRA) is also upregulated in LFABP^−/−^ mice, as is the transcript level of dual-specificity phosphatase 1 (DUSP1); these are known to be induced during adipocyte differentiation [33] and early proliferation of adipocyte progenitors [34], respectively, thus their upregulation may indicate enhanced adipogenic capacity in the LFABP^−/−^ iWAT depot relative to WT. Transcript levels of genes involved in growth arrest, which is a crucial and necessary step during adipogenesis, are also upregulated in HF-fed LFABP null mice, including *Cdkn1a* and *Gadd45g* (Fig. 6, Table S2). Interestingly, preliminary analysis of primary cultures of inguinal APCs isolated from HF-fed WT and LFABP null mice show that fewer APCs differentiated from the LFABP^−/−^ iWAT and those appeared to accumulate bigger lipid droplets, relative to the WT inguinal APCs (Fig. S2). Therefore, the more numerous, smaller adipocytes in the LFABP null mice are likely due to *in vivo* factors*, i.e.*, are not cell autonomous. This is consistent with LFABP being expressed in the liver and the proximal small intestine, but not in AT.

**Figure 6.**
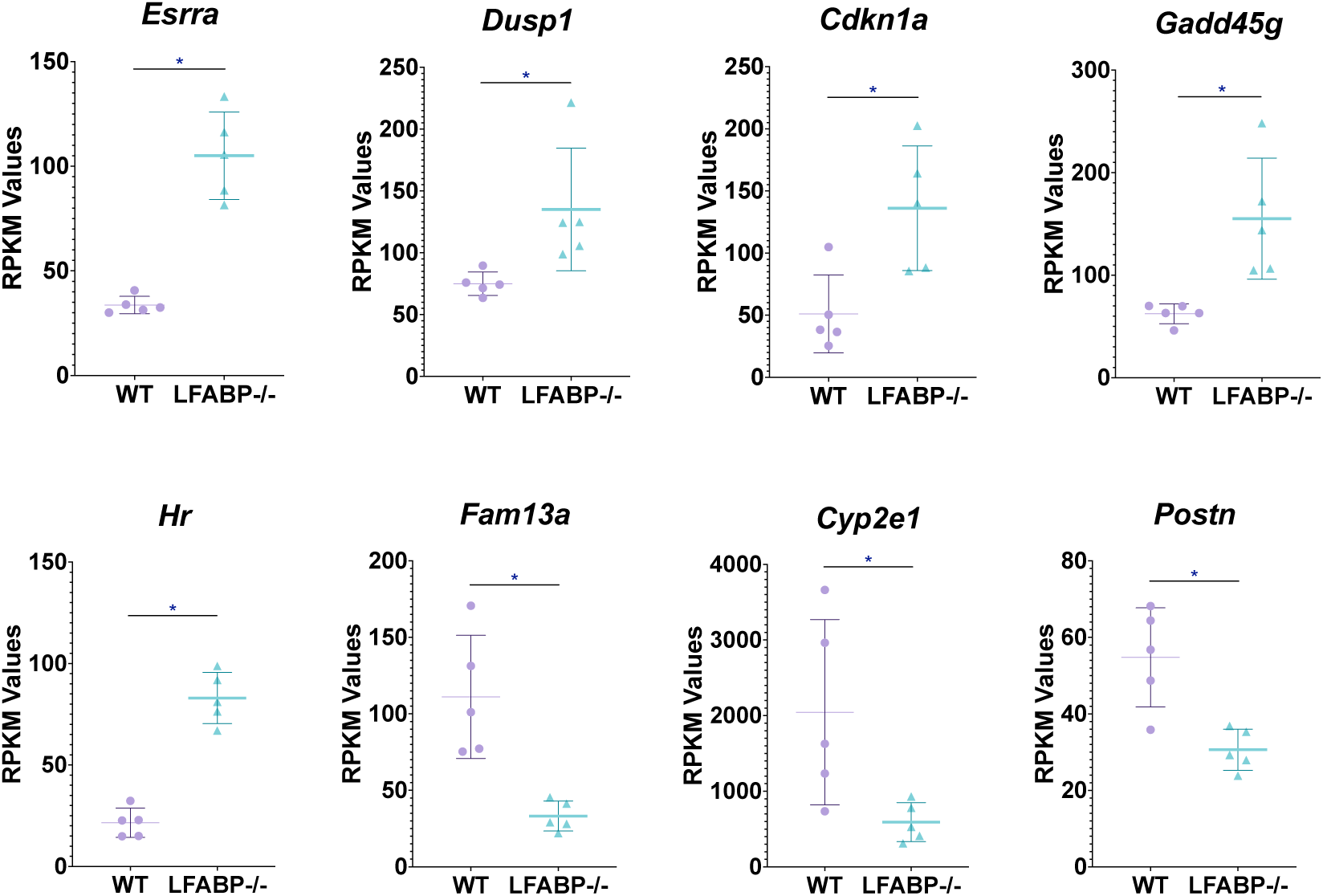
*Lfabp* deletion modifies the expression of genes reportedly related to adipogenesis. RNAseq reveals upregulated transcript levels for *Esrra*, *Insig1*, *Hr*, *Cdkn1a*, *Gadd45g*, and downregulated transcript levels for *Cyp2e1*, *Fam13a*, and *Postn*, involved in cellular processes that may contribute to hyperplastic adipose tissue expansion in the iWAT of HF-fed LFABP null mice relative to the WT (n = 5 per genotype). FDRq < 0.1, |FC| > 1.2, RPKM >20. LFABP*^−/−^ vs.* WT: *\**, *p* < .01. WT, wild-type; LFABP*^−/−^*, liver fatty acid-binding protein null; RPKM, Reads Per Kilobase of transcript per Million reads mapped; Esrra, Estrogen Related Receptor Alpha; Dusp1, Dual-Specificity Phosphatase 1; Cdkn1a, Cyclin-Dependent Kinase Inhibitor 1A; Gadd45g, Growth Arrest and DNA-Damage-Inducible 45 Gamma; Hr, Hairless; Fam13a, Family with Sequence Similarity 13 Member A; Cyp2e1, Cytochrome P450 Family 2 Subfamily E Member 1; Postn, Periostin.

### *Lfabp* deficiency may enhance cholesterol biosynthesis and subsequent mTORC1-mediated iWAT growth

All 3 MSigDBs used for RNA-seq pathway analyses reveal 2 to 3-fold increases in cholesterol biosynthesis and homeostasis pathways in the iWAT of HF-fed LFABP^−/−^ mice (Fig. 4). Indeed, approximately 2 to 2.5-fold increases in the activation of cholesterol biosynthesis and SREBP gene expression pathways are found in the LFABP^−/−^ mouse model upon DIO (FDRq <0.001 for both) (Fig. 7A). Expression levels of 3-hydroxy-3-methylglutaryl-CoA synthase 1 (*Hmgcs1*) (FDRq = 0.036) are increased in the LFABP^−/−^ iWAT, suggesting upregulation of the early steps of cholesterol biosynthesis. Transcript levels of *Insig1* are also upregulated in HF-fed LFABP^−/−^ mice, possibly acting as a brake on cholesterol biosynthesis. Intriguingly, the PI3K/Akt/mTOR signaling pathway is enriched by 33% and that of the protein kinase mechanistic target of rapamycin complex 1 (mTORC1), which is involved in cell growth and cell survival [35] and has been shown to be activated by cholesterol [35], is enriched by 86% in the HF-fed LFABP^−/−^ mice relative to WT (Table 2). This is consistent with evidence demonstrating that increased INSIG1 is rapidly compensated by activation of mTORC1 to restore SREBP1-mediated *de novo* lipogenesis gene expression [36]. Evidence of increased lipid synthesis more generally is seen in the 2.5-fold enrichment in the stearoyl-CoA-desaturase 2 (*Scd2)* transcripts in the HF-fed LFABP^−/−^ relative to WT mice (FDRq = 0.071) (Fig. 7A). Importantly, we found that free cholesterol levels in the iWAT of HF-fed LFABP^−/−^ mice were 50% higher compared to those of WT mice (*P* = 0.003) (Table 1 and Fig. 7B), corroborating the gene expression profiling results.

**Figure 7.**
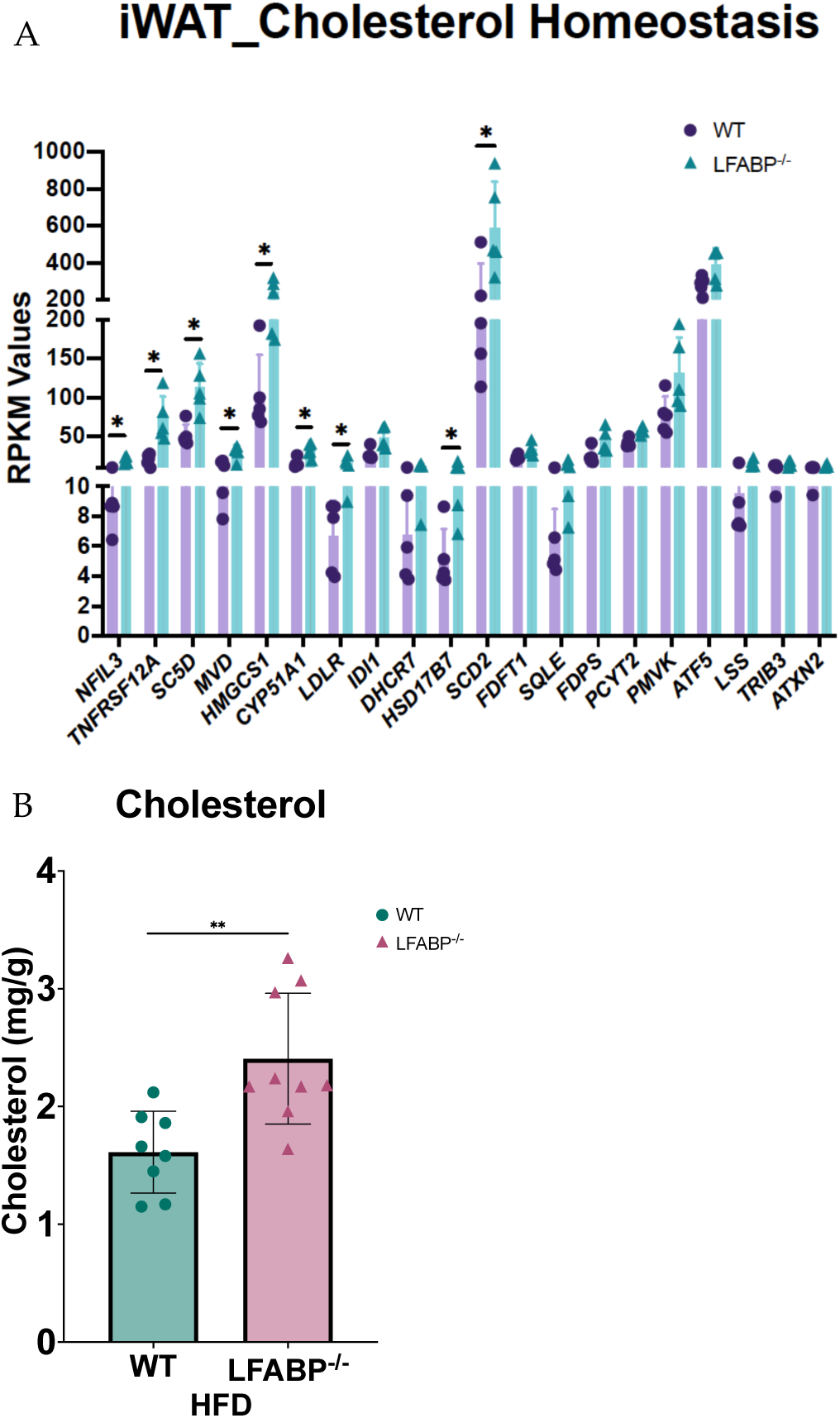
Upregulation of cholesterol metabolism genes and increased cholesterol content of iWAT upon *Lfabp* ablation and HFD challenge. **(A)** Gene expression; n = 5 per genotype. LFABP*^−/−^ vs.* WT: *\**, *p* < .01. **(B)** Free cholesterol content; WT, n = 8; LFABP*^−/−^* n=9. LFABP*^−/−^ vs.* WT: *\*\**, *p* < .001. 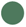 WT, wild-type; 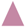 LFABP*^−/−^*, liver fatty acid-binding protein null; HFD, high-fat diet; iWAT, inguinal white adipose tissue.

### *Lfabp* deletion induces iWAT immune and inflammatory responses, and ECM and angiogenesis remodeling

The TNFα signaling pathway via NF-*κ*B, Dectin1-mediated noncanonical NF-*κ*B signaling, and the TNFR2 noncanonical NF-*κ*Β pathway, all involved in cellular immune and inflammatory pathways, are 1.9-(HALLMARK: FDRq <0.001), 1.7-(REACTOME: FDRq = 0.095), and 1.7-fold (REACTOME: FDRq = 0.092) enriched, respectively, in HF-fed LFABP^−/−^ mice, relative to WT. In particular, iWAT transcript levels of TNF receptor superfamily member 12A *(Tnfrsf12a)* are approximately 4-fold higher (FDRq <0.001); TNFRSF12A serves as the specific receptor of the TNF-like weak inducer of apoptosis (TWEAK or TNFSF12A) [37–40] and is involved in non-canonical NF-*κ*B signaling.

Changes in multiple ECM-related pathways were found in the iWAT of HF-fed LFABP^−/−^ mice, including collagen formation and degradation, biosynthesis of collagen and modifying enzymes, assembly of collagen fibrils and other multimeric structures, crosslinking of collagen fibrils, degradation of the ECM, activation of matrix metalloproteinases, ECM proteoglycans, ECM organization, and laminin interactions (Table 2 and S2, Fig. 4B & C).

The transcriptomic profile of iWAT reveals that, among the ECM components, levels of several collagens (*Col1a1*, *Col1a2*, *Col3a1*), along with *Postn* (periostin), *Lama2* (laminin α-2 chain), *Spp1* (osteopontin), and *Ecm1* (extracellular matrix protein 1) are significantly downregulated, whereas *Thbs1* (thrombospondin-1) is significantly upregulated in HF-fed LFABP^−/−^ mice relative to WT (Fig. 8A). The expression level of *Adamts12*, an ECM constructive enzyme, was downregulated in the LFABP^−/−^ mouse model upon DIO, possibly indicating decreased collagen levels due to impaired proteolytic cleavage. The expression levels of degrading enzymes MMP2 (matrix metalloproteinase 2) and MMP9 were also downregulated, as was the level of their inhibitor TIMP2 (tissue metalloproteinase inhibitor 2), presumably explaining the similar transcript levels of the collagen types COL-4 and 5 between LFABP^−/−^ and WT mice (Fig. 8A). By contrast, expression levels of the serine protease inhibitor clade E member 1 (*Serpine1*), also known as plasminogen activator inhibitor-1 (PAI-1), are significantly elevated in the LFABP^−/−^ iWAT, possibly indicating inhibition of the ECM fibrinolytic system and, thus, ECM degradation (Fig. 8A).

**Figure 8.**
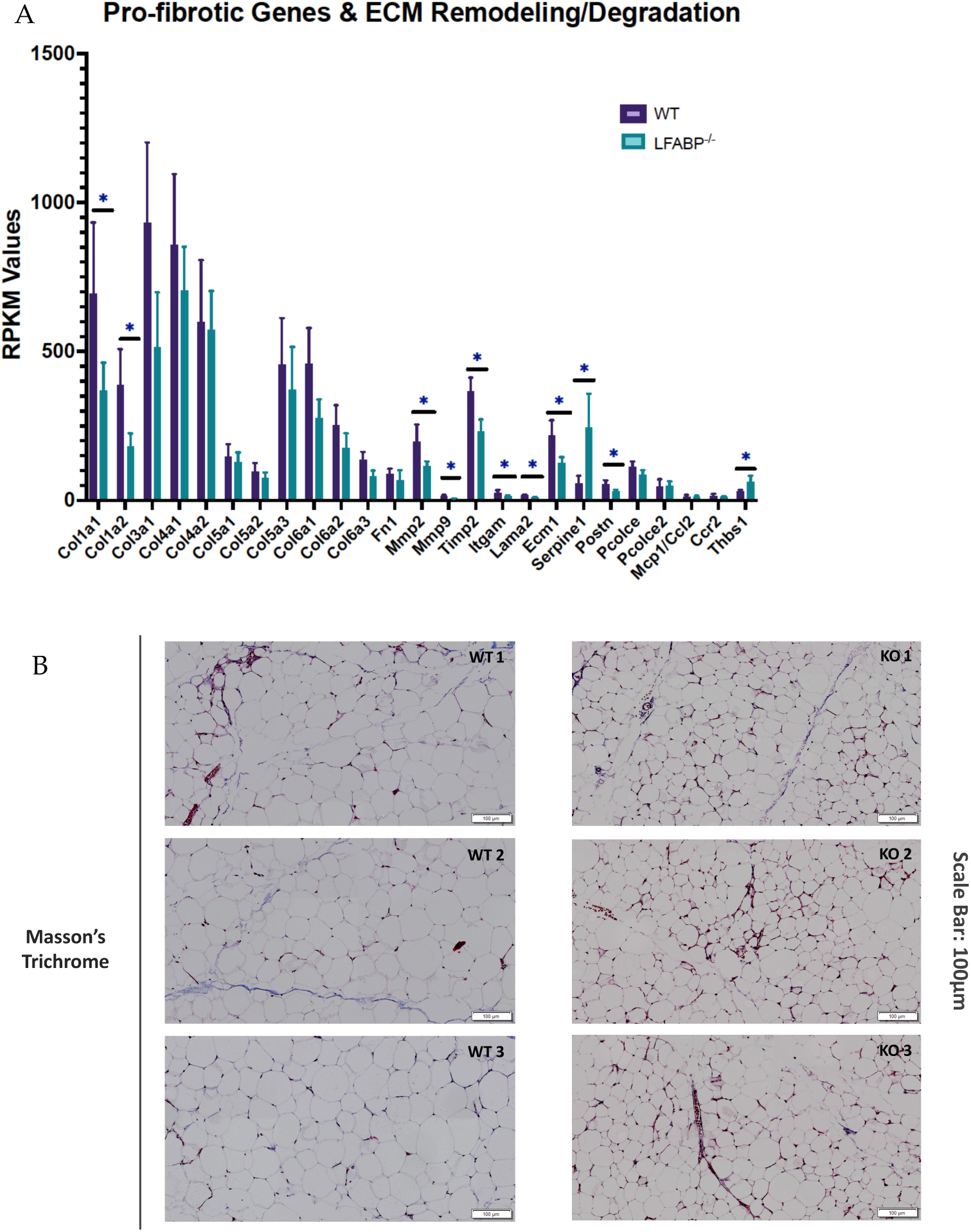
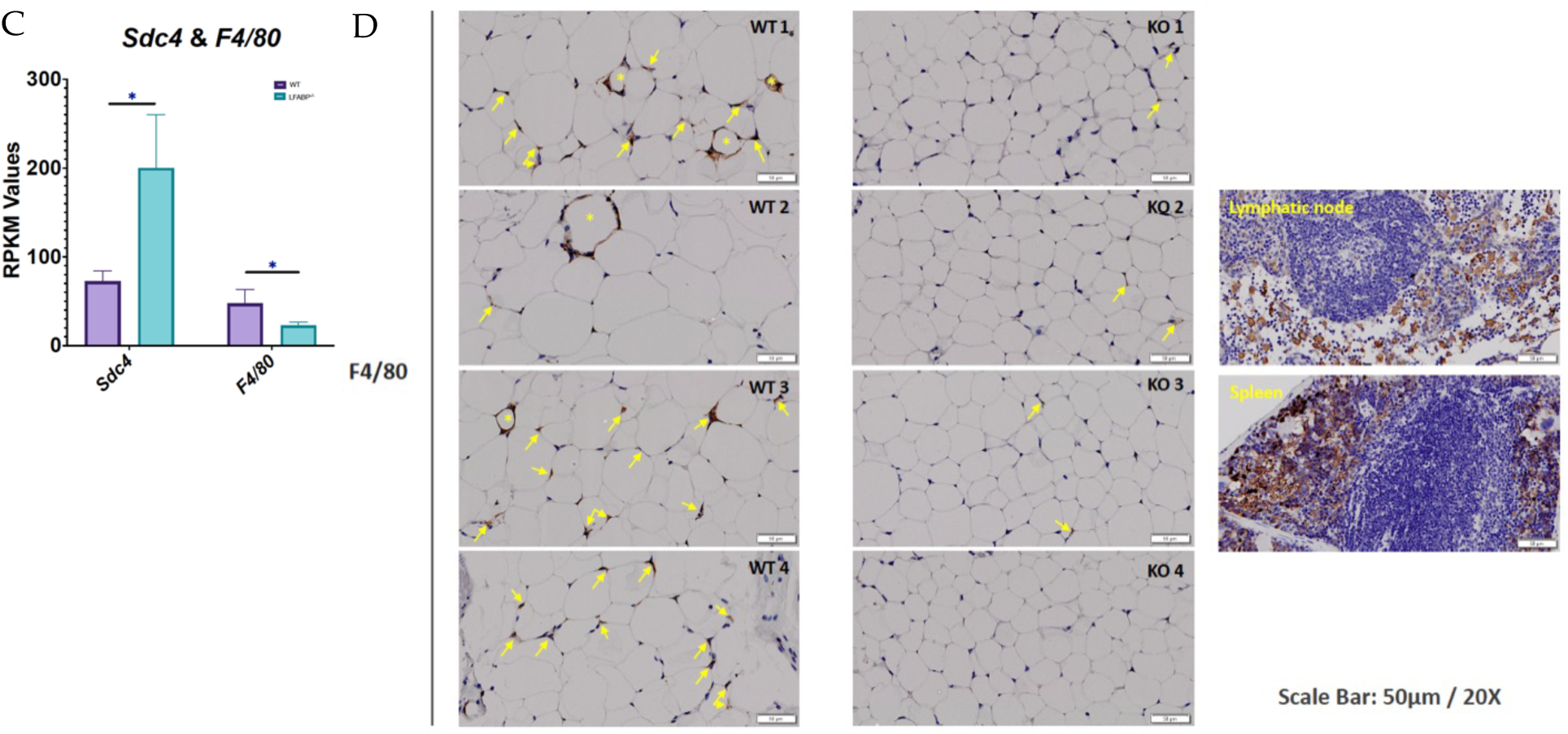
*Lfabp* deletion changes ECM remodeling dynamics, possibly causing adaptive fibrosis and protecting against inflammation in subcutaneous white adipocytes upon DIO. **(A)** RPKM values of pro-fibrotic and ECM remodeling/degrading components in the iWAT of LFABP*^−/−^* mice relative to WT (n = 5 per genotype). **(B)** Representative Masson’s Trichrome staining of iWAT sections in HF-fed WT and LFABP*^−/−^* mice. Magnification: 20x. Scale bar, 100 µm (n = 3 per genotype, from different animals). **(C)** RPKM values of *Sdc4* and *F4/80* in HF-fed LFABP*^−/−^* mice relative to WT (n = 5 per genotype). **(D)** Representative images of IHC staining for F4/80 in the HF-fed LFABP*^−/−^* iWAT compared to the WT. The stars show positive cells organized in CLS around an adipocyte; the arrows show positive cells without specific organization. Positive control for mesenteric lymphatic node and spleen specimens. Magnification: 20x. Scale bar, 50 µm (n = 4 per genotype, from different animals). LFABP*^−/−^ vs.* WT: *\**, *p* < .01; *\*\**, *p* < .001. WT, wild-type; LFABP*^−/−^*, liver fatty acid-binding protein null; KO, knockout; Sdc4, Syndecan 4; F4/80/Adgre1, Adhesion G Protein-Coupled Receptor E1; CLS, Crown-like structure; RPKM, Reads Per Kilobase of transcript per Million reads mapped. The results should describe the experiments performed and the findings observed. The results section should be divided into subsections to delineate different experimental themes. Subheadings should be descriptive phrases. All data must be shown either in the main text or in the Supplementary Materials.

SERPINE1 has been shown to induce cell proliferation and migration, as well as pro-angiogenic activity [41–44]. Indeed, RNA-seq analysis for a set of angiogenesis-related transcripts shows similar expression levels between the knockout (KO) and WT animals, suggesting preserved angiogenesis in the iWAT of HF-fed LFABP^−/−^ mice (Fig. S8A). Validation of RNA-seq results with targeted qRT-PCR for selected angiogenesis-related genes shows upregulated mRNA levels of *Vegf*, its receptor *Vegfr2*, and *Nos2* in the iWAT of HF-fed LFABP null mice relative to the WT (Fig. S8b). Moreover, Masson’s trichrome staining for fibrosis shows no appreciable differences between the HF-fed LFABP null and WT mice. It is of interest, however, that small adipocytes are found close to more fibrotic areas in the KO iWAT sections, which is consistent with the concepts of ‘adaptive fibrosis’ and ‘homeostatic adipogenesis’ [45–47] (Fig. 8B).

The transcript levels of *Sdc4* (syndecan 4), involved in ECM interactions and vasculogenesis, are increased, whereas those of *F4/80* (or *Adgre1*), characteristic of macrophage infiltration, are decreased in the HF-fed LFABP^−/−^ iWAT relative to WT (Fig. 8C). To directly assess macrophage infiltration, iWAT sections were stained for F4/80; in keeping with the gene expression data, markedly decreased infiltration of macrophages in the subcutaneous adipose tissue of the MHO LFABP ablated mice relative to the WT was observed, including large areas with no apparent macrophage staining at all. Those positive cells found in the iWAT of LFABP^−/−^ mice are mostly single, mildly stained isolated macrophages. In contrast, in the iWAT of WT mice, there were more areas showing the presence of macrophages, and the greater intensity of the staining indicates the aggregation of more cells compared to the iWAT of the KOs. Notably, whereas in the iWAT of HF-fed WT mice we detected areas with positive cells organized in CLSs around adipocytes, CLSs were not found in the iWAT of the LFABP^−/−^ mice (Fig. 8D), in keeping with the MHO phenotype of these mice.

Collectively, these results reveal that while the WT responds as expected to HFD by developing inflamed adipose tissue with increased pericellular fibrosis, the LFABP KO manages to prevent this, showing neither inflammation nor macrophage infiltration in iWAT upon DIO.

## 4. Discussion

### Evidence of hyperplastic WAT expansion upon *Lfabp* deletion and HF-feeding

Subcutaneous adipose tissue is the largest and at the same time the least metabolically harmful AT depot in the body [15]. With the development of obesity, WAT undergoes a process of tissue remodeling in which adipocytes increase in both number and size [16, 23, 48, 49]. Evidence suggests that the recruitment of APCs is a feature of healthy AT expansion to meet the need of storing excess energy. This process necessitates remodeling of the fibrous ECM [11, 50], which may become a limiting factor for adipocyte size in the context of obesity [11].

LF-fed LFABP^−/−^ mice have higher fat mass compared to WT mice, with the size of white adipocytes similar under LF-feeding. Remarkably, though, whereas the WT mice respond as expected when challenged with HFD, the LFABP^−/−^ iWAT depot becomes populated by approximately 5-fold higher numbers of inguinal adipocytes, whose size is only 25% that of their WT counterparts. This strongly suggests an unusual hyperplastic expansion of subcutaneous fat in the LFABP^−/−^ mice. Smaller adipocytes are thought to be more insulin sensitive [9, 16, 48, 51], thus the effect of *Lfabp* ablation on the quality of sWAT may contribute to the previously reported MHO phenotype of the LFABP^−/−^ mouse, which is characterized by normal plasma glucose and insulin levels despite marked obesity [3][3].

### *Lfabp* deletion induces alterations in the transcriptomic profile of subcutaneous fat suggestive of an enhanced adipogenic program

The significantly upregulated expression of several pathways, particularly those of cholesterol metabolism and biosynthesis, mTORC1, and PI3K/Akt/mTOR signaling, are indicative of enhanced tissue growth in HF-fed LFABP^−/−^ mice relative to WT. Moreover, the modulation of specific transcripts in the iWAT of HF-fed LFABP null mice relative to WT point to healthy AT expansion. For example, the decreased expression of *Cyp2e1* is of particular interest, as its expression is enriched in CD142^+^ cells, a subpopulation of adipose stem and progenitor cells (ASPCs) that has been attributed non- and anti-adipogenic properties both *in vivo* and *in vitro* [30, 52]. *Cyp2e1*-deficient mice, moreover, were shown to be protected against DIO and insulin resistance [53]. Thus, the robust downregulation of *Cyp2e1* in the iWAT of LFABP^−/−^ mice may be indicative of enhanced adipogenic capacity and improved glycemic control, as found in these mice [4].

Lower levels of *Fam13a* were also found in the hyperplastic iWAT of HF-fed LFABP^−/−^ mice. A murine model of *Fam13a* deficiency exhibited higher numbers of small adipocytes in iWAT, enhanced adipogenic potential, and preserved glucose uptake and insulin responsiveness [31], findings that are very similar to those observed herein in HF-fed LFABP^−/−^ mice, suggesting that *Fam13a* downregulation may contribute to the hyperplastic iWAT phenotype in these mice.

The iWAT of HF-fed LFABP null mice was also found to have enriched *Esrra* transcripts relative to the WT. ESRRA deficiency in mice is characterized by reduced adiposity, possibly due to a decrease in APCs differentiation and adipogenesis in WAT [33, 54, 55]. Thus, enriched *Esrra* transcript levels herein further support the observed increased cellularity and enhanced adipogenic potential in the HF-fed LFABP^−/−^ iWAT. Interestingly, *Esrra* has been additionally shown to regulate transcription of genes participating in autophagy [56], which is progressively induced during adipocyte differentiation such that loss of autophagy may compromise white adipogenesis [57–60]. Cholesterol has been shown to serve as an endogenous ligand for ESRRA [56, 61–64], thus the significant increases found in cholesterol biosynthesis pathways and iWAT cholesterol levels in HF-fed LFABP^−/−^ mice may be related to increased *Esrra* -mediated adipogenesis.

Transcript levels of *Scd2* were found in the LFABP^−/−^ mouse model. SCD’s are well known to be induced during differentiation of 3T3-L1 preadipocytes into adipocytes and in AT with obesogenic diets [65, 66]. The monounsaturated fatty acids (MUFAs), products of SCDs, are critical components of cellular membrane phospholipids (PLs), cholesterol esters, and TAG stores, contributing to appropriate adipocyte membrane structure and function as well as sufficient capacity for excess energy storage, both of which would be necessary for cell proliferation and growth in newly emerging APCs in the obese LFABP^−/−^ mouse model. Overall, the transcriptomic changes found in iWAT largely support the observed hyperplastic growth of LFABP^−/−^ adipose tissue during HFD feeding.

### Enrichment in cholesterol-related pathways and increased cell cholesterol content may contribute to small, insulin sensitive adipocytes in the iWAT of HF-fed LFABP^−/−^ mice

The transcriptomic data showing a significant upregulation of cholesterol biosynthesis in the HF-fed LFABP^−/−^ mice, relative to the WT, are consistent with the observed higher tissue cholesterol levels. Interestingly, both cholesterol tissue levels and its subcellular distribution are proposed to be involved in adipocyte cell size regulation, where depletion of cholesterol from the plasma membrane has been reported to reproduce insulin resistance-related defects found in enlarged fat cells, possibly via internalization of surface caveolins [46, 67, 68]. Increased sterols combined with the markedly smaller inguinal adipocyte size may suggest increased cholesterol in the plasma membrane relative to the lipid droplets [46], consistent with the maintained insulin sensitivity described previously in the MHO LFABP^−/−^ mouse model [3, 4].

The increased levels of *Insig1* expression in the iWAT of HF-fed LFABP^−/−^ mice is consistent with elevated adipose tissue levels reported at the onset of DIO [69]. Interestingly, it has been shown that expression of *Insig1* progressively increases during maturation of adipocyte progenitors and impedes lipogenesis in mature adipocytes, thereby determining adipocyte size and storage capacity [70]. It has also been reported that INSIG1-mediated blockade of adipose tissue lipogenesis is immediately compensated by activation of mTORC1 to restore SREBP1-dependent *de novo* lipogenesis gene expression [70]. This too is in accordance with the observed upregulation of the PI3K/Akt/mTOR and mTORC1 signaling pathways, as well as increases in the activation of gene expression by SREBP and cholesterol biosynthesis in the LFABP^−/−^ mouse model upon DIO. Cholesterol in turn can mediate activation of the mTORC1 signaling pathway and its downstream anabolic effects, including cell growth, proliferation, metabolism, survival, and angiogenesis [35, 71, 72], as observed herein.

### *Lfabp* ablation alters immune and inflammatory responses in inguinal adipocytes upon DIO

Non-canonical NF-*κ*B signaling can be stimulated after lipopolysaccharide (LPS) binding to Dectin1 and TNFR2 and can be initiated by TNFR2 and TNFRSFs (TNFR superfamily members) such as TNFRSF12a, which is significantly enriched in the iWAT of HF-fed LFABP^−/−^ mice. Overexpression of TNFRSF12A has been shown to activate the non-canonical NF-*κ*B pathway independent of binding to its ligand TWEAK [73], and indeed *Tweak* transcript levels are similar between the LFABP^−/−^ and WT mice. Bacterial LPS-induced inflammation may induce TNFRSF12A expression via TNFα signaling [74], thus the observed *Tnfrsf12* overexpression may be induced by an extracellular stimulus, *e.g.* LPS derived from obesity-related gut bacterial species [75]; we have shown that LPS biosynthetic pathways are significantly upregulated in LFABP^−/−^ intestinal mucosa [76]. In addition to TNFR2, Dectin1, which has been shown to be involved in adaptive immunity and tolerance [77], is also significantly upregulated in the LFABP null mice.

It has been suggested that the inflammatory responses classically attributed to TNFα, including LPS-induced cytotoxicity, are mediated by TNFR1, whereas TNFR2 has been implicated in the suppression of TNFα-induced inflammatory responses and NF-*κ*B-dependent gene expression [78]. It is therefore possible that, despite their adiposity, LFABP^−/−^ mice may be, at least partially, protected against adverse inflammatory responses through non-canonical activation of NF-*κ*B signaling through TNFR2 and Dectin1.

### *Lfabp* deletion may alter inguinal fat ECM remodeling, fibrosis, inflammation, and angiogenesis upon DIO

A biphasic development of the ECM has been suggested to occur during adipogenesis, in which the fibrillar collagen types 1 and 3 are decreased early on and return to their initial levels at the later stages of differentiation [79–81]. In this study, HF-fed LFABP^−/−^ mice display downregulated expression of *Col1a1*, *Col1a2*, and *Col3a1* in iWAT, possibly indicative of newly differentiated adipocytes that have not fully acquired the mature adipocyte phenotype. Intriguingly, *Postn* deficiency has been reported to attenuate LPS-/HFD-induced AT fibrosis and to improve IR, thus the lower *Postn* levels found herein may contribute to healthy hyperplastic AT expansion and maintained insulin sensitivity during obesity [82, 83], as we observe in the LFABP^−/−^ mouse model.

*Spp1* encodes for osteopontin, an ECM protein important in cell adhesion, migration, and ECM degradation [84], which has been demonstrated to inhibit adipogenic and promote osteogenic differentiation in mesenchymal stem cells (MSCs) [85]. Here we found transcriptional downregulation of *Spp1* in the iWAT of HF-fed LFABP^−/−^ mouse. Deficiency of osteopontin results in increased ratios of both subcutaneous and visceral AT to body weight [85], as is found in the LFABP^−/−^ mouse.

We also found increased *Sdc4* transcript levels in the LFABP^−/−^ iWAT, compared to the WT, further corroborating ECM remodeling that may influence adipose tissue remodeling during obesity development. During DIO, *Sdc4* deletion contributes to dyslipidemia, hyperglycemia and insulin resistance, as well as increased adipocyte size and macrophage infiltration [67]. Our findings in HF-fed LFABP^−/−^ mice, which overexpress *Sdc4,* are consistent with these results, since we find a marked decrease in the size of inguinal adipocytes, decreased expression and staining of F4/80, indicative of lower macrophage infiltration, and normal glycemic and lipidemic profiles despite massive adiposity, as previously reported [3]. This too confers additional support for low inflammation and maintained insulin sensitivity in the HF-fed LFABP mouse model.

Elevation of *Serpine1* in the LFABP^−/−^ iWAT may indicate inhibition of the ECM fibrinolytic system and, thus, ECM degradation. While the role of SERPINE1 in adipogenesis and angiogenesis is controversial [68, 86–89], it has been shown that overexpression of SERPINE1 in AT results in larger fat pads with higher adipocyte density[86][86], consistent with the phenotype found in the LFABP^−/−^ iWAT. Additionally, the inhibitory effect of *Serpine1* on ECM degradation could, potentially, serve as a brake on adipocyte size. Indeed, excessive ECM deposition in AT during obesity development has been suggested to contribute to a state of adaptive fibrosis, in which a more rigid ECM may prevent excessive enlargement of adipocytes and preserve adipocyte function [45, 47]. In this regard, APCs have been demonstrated to accumulate in fibrotic areas within the WAT of obese individuals [90]. HF-fed LFABP^−/−^ iWAT stained for Masson’s trichrome show that the numerous small adipocytes described here appear to be aggregated in regions with more intense staining. Scarce macrophage infiltration and negligible or absent CLS formation in the subcutaneous fat of the LFABP^−/−^ mice indicates a lower inflammatory state and maintained metabolic function within the depot, supporting the notion of an ‘adaptive fibrosis’ and a metabolically healthy obese state.

Overall, given the massive iWAT adiposity of the LFABP^−/−^ mouse model, we speculate that their inguinal adipocytes reach a critical/maximal size after which hyperplastic growth is triggered earlier during the HF-feeding period, relative to WT. This would lead to APC recruitment that in turn would regulate ECM remodeling to drive hyperplastic iWAT expansion. It has been reported that hypertrophy precedes hyperplasia/adipogenesis, and that adipocyte hypertrophy and turnover increase depending on the rate of change of overall fat mass [9, 91]. Therefore, the fact that LFABP^−/−^ mice challenged with HF-feeding become more obese compared with their WT counterparts, in combination with decreased ECM degradation that would render ECM in iWAT more rigid, may offer a reasonable explanation for the presence of smaller adipocytes. Notably, the phenotype is different on LF-feeding, where *Lfabp* ablation appears to predispose for higher iWAT mass relative to the WT, despite similar BW, and where adipocyte size is comparable to WT.

While the SAT hyperplastic phenotype is clear, it is worth noting that only male mice were studied in depth thus far. Additionally, subcutaneous WAT was thoroughly examined, whereas visceral WAT and interscapular BAT were not extensively studied apart from cellularity and histological features. Given the dramatic changes observed in BAT, particularly, future studies will characterize transcriptional and physiological changes in BAT, which may contribute to the MHO phenotype of the obese LFABP null mice. It will also be informative to examine the time course of adipogenesis *in vivo*, and to further assess transcriptomic changes using single cell RNA-seq to dissect mechanistic pathways at the adipocyte level that contribute to DIO-dependent hyperplastic expansion in the LFABP^−/−^ mouse.

## 5. Conclusions

In summary, we show that while WT animals respond as expected to DIO by developing hypertrophied inguinal adipocytes and inflamed iWAT with increased pericellular fibrosis, the LFABP null mouse model, by contrast, expands by increasing adipocyte number, not size, and is protected from inflammation and macrophage infiltration (Fig. 9). Since LFABP is not expressed in AT, these results suggest that its deficiency promotes interorgan signaling that affects adipocyte cellularity, limiting hypertrophy and driving hyperplasia in expansion of potentially metabolically beneficial subcutaneous iWAT. The previously described MHO phenotype of LFABP null mice, characterized by normal glucose and insulin levels as well as decreased hepatic steatosis [3] and resistance to obesity-induced decrease in exercise capacity [4], is further corroborated in the present study, where we show a highly unusual hyperplastic mode of subcutaneous fat expansion. The mechanisms by which LFABP, a lipid transport protein expressed within the liver and the proximal intestine, impacts AT and, as we showed previously, muscle tissue [4], remains an intriguing question. We found increased gut microbial diversity [75] and intestinal mucosal anandamide (AEA) levels [3] upon LFABP ablation and have recently observed significantly smaller gall bladder size in the HF-fed LFABP null mouse, relative to WT, as well as alterations in plasma BA levels and composition [76]. Since LFABP binds both bile acids and endocannabinoids, we speculate that bile acid-gut microbiota metabolism and endocannabinoid system-related signaling may be involved in the effects of LFABP ablation on peripheral tissues and systemic metabolism.

**Figure 9.**
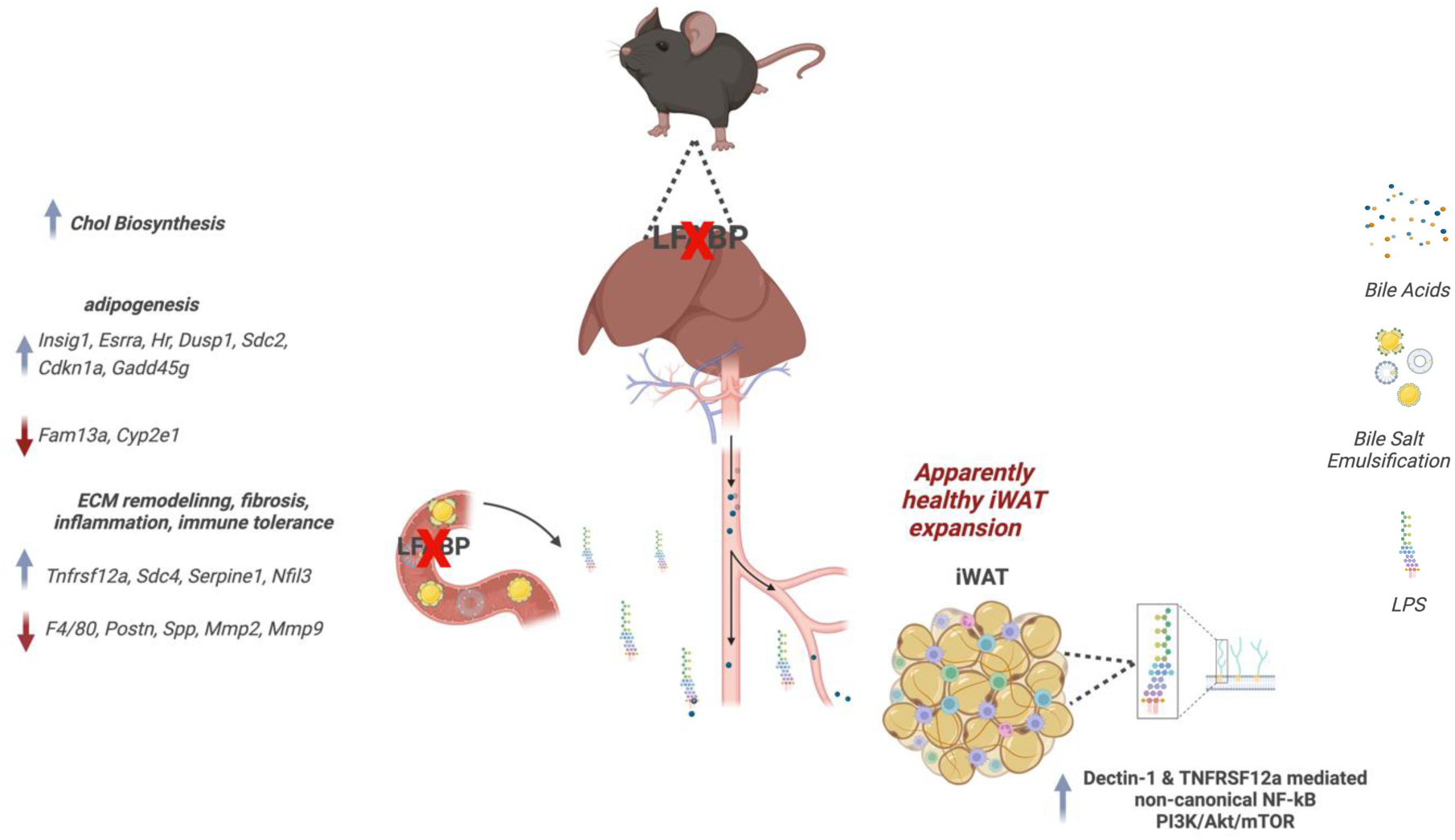
*Lfabp* deletion and HF-feeding lead to hyperplastic expansion of iWAT. Whole-body *Lfabp* ablation results in heavier iWAT depot with more numerous and smaller adipocytes, *i.e.,* hyperplastic iWAT expansion. Transcriptomic modifications of genes reportedly involved in enhanced adipogenesis, as well as ECM remodeling, inflammation, and immune tolerance are suggested to participate in the observed adipocyte phenotype. Additionally, upregulation of pathways involved in cholesterol biosynthesis, non-canonical NF-*κ*B signaling upon potential LPS-mediated Dectin-1 and TNFRSF12a stimulation, and PI3K/AKT/mTOR signaling are also proposed to contribute to hyperplastic iWAT growth upon *Lfabp* deletion and DIO. Chol, cholesterol; ECM, extracellular matrix; iWAT, inguinal white adipose tissue; LFABP, liver fatty acid-binding protein; LPS, lipopolysaccharide; TNFRSF12a, Tumor Necrosis Factor Receptor Superfamily Member 12A.

## Supplementary Materials

The following supporting information can be downloaded at: www.mdpi.com/xxx/s1, Figure S1: iWAT RNAseq Volcano Plot; Figure S2: Primary culture of stromal vascular fraction (SVF) isolated from the iWAT of HF-fed LFABP null and WT male mice; Table S1: Diet compositions of LF and HF diets; Table S2: HALLMARK, KEGG and REACTOME Gene Sets Enriched in the iWAT of HF-fed LFABP^−/−^ male mice; Table S3: HALLMARK, KEGG and REACTOME Gene Sets Downregulated in the iWAT of HF-fed LFABP-/- male mice.

## Author Contributions

Conceptualization: A.D., J.S.; Data curation: S.H., A.D.; Formal Analysis: A.D., A.F., S.H.; Funding Acquisition: J.S., S.K.F.; Investigation: A.D., Y.Z., J.S., A.F.; Methodology: A.D., J.S., S.K.F.; Project administration: J.S., A.D.; Resources: A.D., J.S., Y.Z.; Software: S.H., A.D.; Supervision: J.S., S.K.F., L.S.S.; Validation: A.D., J.S., S.K.F.; Visualization: A.D., J.S.; Writing—original draft: A.D.; Writing—review & editing: J.S., S.K.F. All authors have read and agreed to the published version of the manuscript.

## Funding

This work was supported by a National Institutes of Health Grant DK-38389 (JS), Funds from the New Jersey Agricultural Experiment Station NJ14163 (JS), and the National Institutes of Health, Institute of Diabetes and Digestion and Kidney Diseases (NIDDK) (USA) Grant R01 DK121547 (SKF).

## Institutional Review Board Statement

The animal study protocol was approved by the Rutgers University Institutional Animal Care and Use Committee (IACUC) (PROTO999900318 18 June 2023).

## Data Availability Statement

All data needed to evaluate the conclusions in the paper are present in the paper and/or the Supplementary Materials. Further information and requests for resources and reagents should be directed to the lead contacts JS and AD. RNA-Seq data have been deposited to the GEO (Gene Expression Omnibus database), with the GEO Series number: GSE277001.

## Acknowledgments

The authors thank Dr. Marianne Polunas of the Rutgers University Research Pathology Services for invaluable help with tissue histology. A special thanks to Dr. Kalypso Karastergiou for her invaluable help and advice on RNA-seq analysis using GSEA.

## Conflicts of Interest

The authors declare no conflicts of interest. The funders had no role in the design of the study; in the collection, analyses, or interpretation of data; in the writing of the manuscript; or in the decision to publish the results.

## Abbreviations

The following abbreviations are used in this manuscript:

LFABP^−/−^: Liver Fatty Acid-Binding Protein Null
HFD: High-Fat Diet
SAT: Subcutaneous Adipose Tissue
WT: Wild-Type
LCFAs: Long-Chain Fatty Acids
MAGs: Monoacylglycerols
eCBs: Endocannabinoids
BAs: Bile Acids
MHO: Metabolically Healthy Obese
FFA: Free Fatty Acid
AT: Adipose Tissue
APCs: Adipocyte Progenitor Cells
ECM: Extracellular Matrix
IR: Insulin Resistance
sWAT: Subcutaneous White Adipose Tissue
vWAT: Visceral WAT
BAT: Brown Adipose Tissue
DIO: Diet-Induced Obesity
LFD: Low-Fat Diet
MRI: Magnetic Resonance Imaging
H&E: Hematoxylin and Eosin
CLSs: Crown-Like Structures
FCV: Fat Cell Volume
FCN: Fat Cell Number
TAG: Triacylglycerol
iWAT: Inguinal WAT
eWAT: Epididymal
WAT iBAT: Interscapular BAT
GSEA: Gene Set Enrichment Analysis
MSigDB: Metabolic Signatures Database
NES: Normalized Enrichment Score
FDR: False Discovery Rate
RPKM: Reads Per Kilobase per Million mapped
PBS: Phosphate-Buffered Saline
BW: Body Weight
CYP2E1: Cytochrome P450 family 2 subfamily E member 1
Areg: Adipogenesis Regulator
FAM13A: Family with sequence similarity 13 member A
Hr: Hairless
ESRRA: Estrogen-Related Receptor α
DUSP1: Dual-Specificity Phosphatase 1
Cdkn1a: Cyclin-Dependent Kinase Inhibitor 1A
Gadd45g: Growth Arrest and DNA-Damage-Inducible 45 Gamma
Postn: Periostin
Hmgcs1: 3-hydroxy-3-methylglutaryl-CoA synthase 1
mTORC1: Mechanistic Target of Rapamycin Complex 1
Scd2: Stearoyl-CoA-Desaturase 2
Tnfrsf12a: Tumor Necrosis Factor Receptor Superfamily Member 12A
TWEAK: TNF-like weak inducer of apoptosis
Col: Collagen
Lama2: Laminin α-2 chain
Spp1: Osteopontin
Ecm1: Extracellular Matrix Protein 1
Thbs1: Thrombospondin-1
MMP: Matrix Metalloproteinase
TIMP2: Tissue Metalloproteinase Inhibitor 2
Serpine1: Serine Protease Inhibitor Clade E Member 1
PAI-1: Plasminogen Activator Inhibitor-1
KO: Knockout
Sdc4: Syndecan 4
ASPCs: Adipose Stem and Progenitor Cells
MUFAs: Monounsaturated Fatty Acids
PLs: Phospholipids
LPS: Lipopolysaccharide
TNFRSFs: NFR Superfamily Members
MSCs: Mesenchymal Stem Cells
AEA: Anandamide
Chol: Cholesterol

## Notes

https://app.globus.org/groups/b4841dd0-f8ea-11ee-a017-1fb3dd6261f9/about

https://app.globus.org/file-manager?origin_id=d372c739-3e77-4e37-bc09-afd725d6be14&origin_path=%2FStorch%2FStorch_30RNA_210809%2FFastq%2F

### Competing Interest Statement

The authors have declared no competing interest.

