## Supplemental Material for "Novel hyperplastic expansion of white adipose tissue underlies the metabolically healthy obese phenotype of male LFABP null mice"

Anastasia Diolintzi *et al.*

**This PDF file includes:**

Figs. S1 to S2

Tables S1 to S3

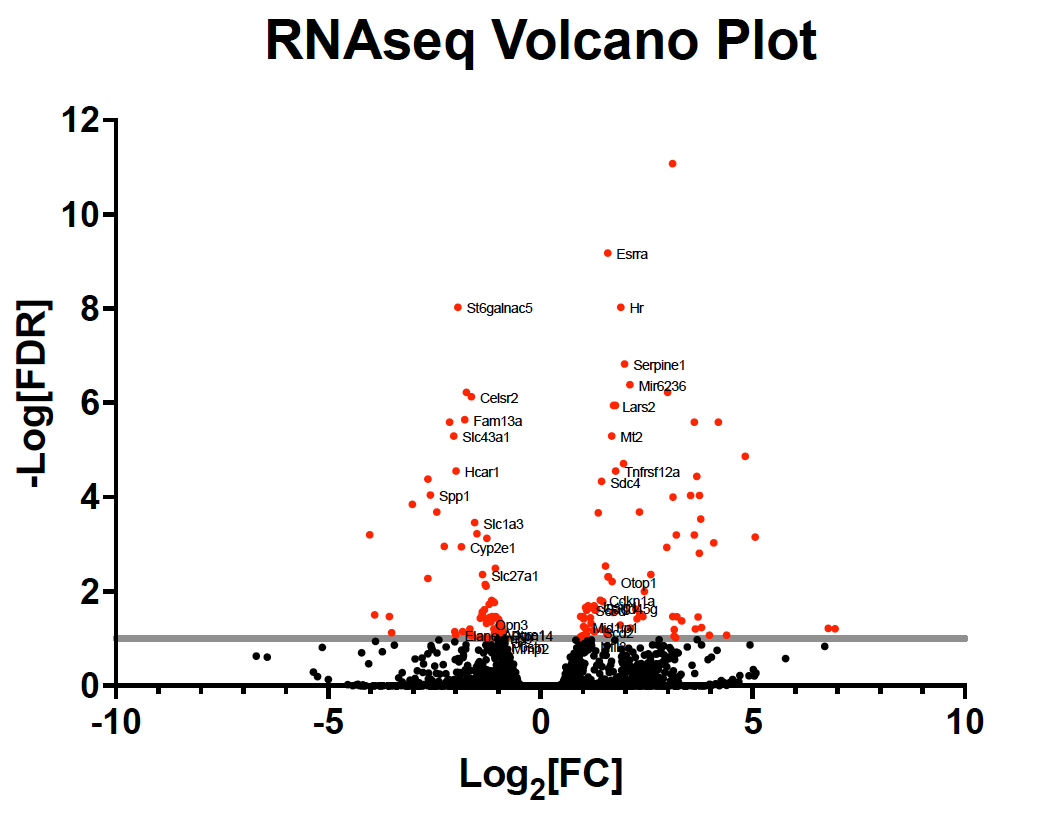

**Fig. S1. iWAT RNAseq Volcano Plot.** Volcano plot of RNAseq analysis for iWAT between LFABP null and WT mice (n = 5 / genotype). The X axis shows the logarithmic value with base 2 for transcript fold change (Log_2_[FC]). The Y axis shows the negative logarithmic value of FDRq-values (-Log[FDR]). The solid gray line indicates the -Log[FDR] for FDRq-value < 0.1, *i.e.,* 1. Selected transcript names are shown for some data points.

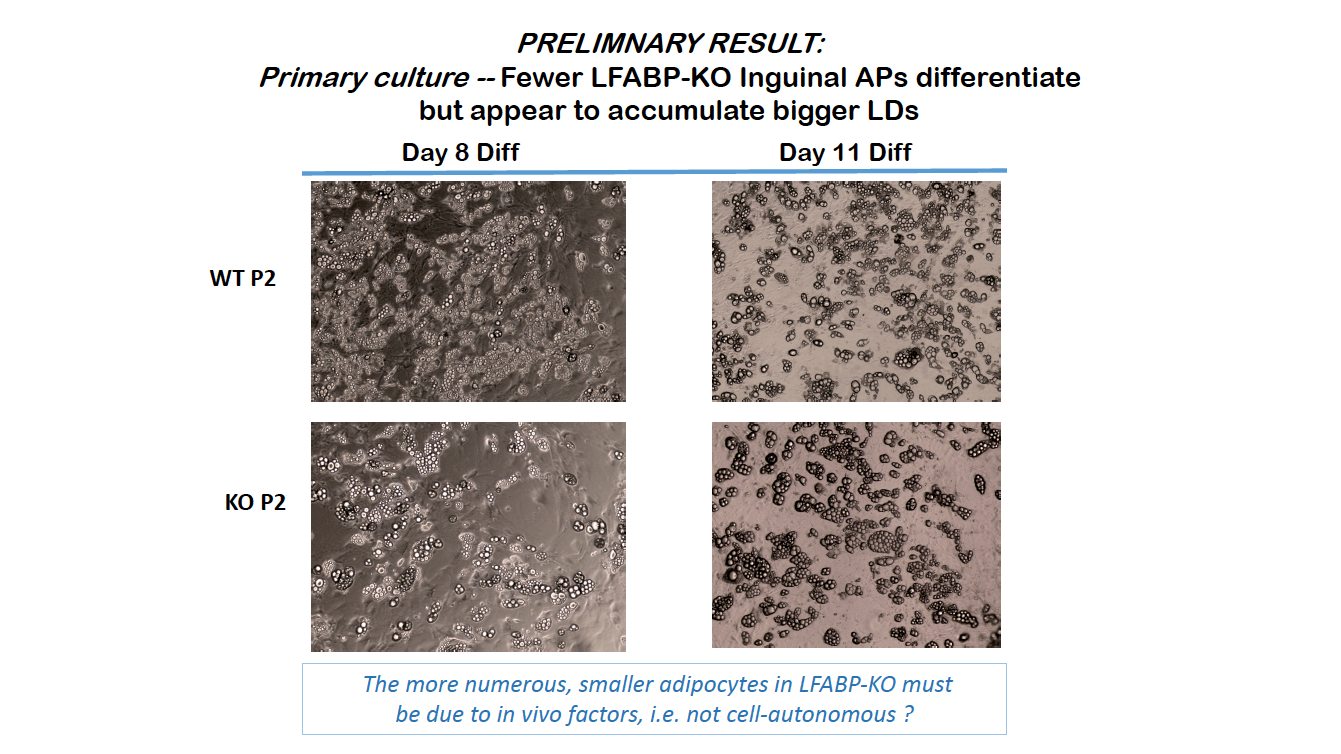

**Fig. S2. Primary culture of stromal vascular fraction (SVF) isolated from the iWAT of HF-fed LFABP null and WT male mice.** Fewer LFABP null inguinal APCs differentiate but appear to accumulate bigger lipid droplets. The more numerous, smaller adipocytes in LFABP null mice must be due to *in vivo* factors, *i.e.*, not cell autonomous. For each genotype pooled SVF from the iWAT of two mice was used for cell culture experiments. WT, wild type; KO, knockout; P, passage

|  | **LFD** | | | **HFD** | | |
| --- | --- | --- | --- | --- | --- | --- |
|  | *g* | *Kcal* | *% Kcal* | *g* | *Kcal* | *% Kcal* |
| Casein | 200 | 800 | 19.7 | 200 | 800 | 19.7 |
| L-Cystine | 3 | 12 | 0.3 | 3 | 12 | 0.3 |
| Corn starch | 315 | 1260 | 31 | 72.8 | 291 | 7.2 |
| Maltodextrin | 35 | 140 | 3.5 | 100 | 400 | 9.9 |
| Sucrose | 350 | 1400 | 34.5 | 172.8 | 691 | 17 |
| Cellulose | 50 | 0 | 0 | 50 | 0 | 0 |
| Soybean oil | 10 | 90 | 2.2 | 10 | 90 | 2.2 |
| Lard | 8.5 | 77 | 1.9 | 0 | 0 | 0 |
| Cocoa butter | 26.5 | 239 | 5.9 | 192.5 | 1733 | 42.7 |
| High oleic safflower oil | 0 | 0 | 0 | 0 | 0 | 0 |
| Mineral mix, S10026 | 10 | 0 | 0 | 10 | 0 | 0 |
| Dicalcium phosphate | 13 | 0 | 0 | 13 | 0 | 0 |
| Calcium carbonate | 5.5 | 0 | 0 | 5.5 | 0 | 0 |
| Potassium citrate, 1 H_2_O | 16.5 | 0 | 0 | 16.5 | 0 | 0 |
| Vitamin mix, V10001 | 10 | 40 | 1 | 10 | 40 | 1 |
| Choline bitartrate | 2 | 0 | 0 | 2 | 0 | 0 |
| FD&C yellow dye no. 5 | 0.05 | 0 | 0 | 0 | 0 | 0 |
| FD&C yellow dye no. 40 | 0 | 0 | 0 | 0.05 | 0 | 0 |
| FD&C yellow dye no. 1 | 0 | 0 | 0 | 0 | 0 | 0 |
| Total | 1055.05 | 4057 | 100 | 858.15 | 4057 | 100 |

Table S1. Diet compositions of LF and HF diets. Adjusted from (*1*)

| **Gene Set** | **NES** | **FDR** |
| --- | --- | --- |
| REACTOME_Cholesterol Biosynthesis | 2.52 | 0.000 |
| HALLMARK_Cholesterol Homeostasis | 2.28 | 0.000 |
| REACTOME_Activation of Gene Expression by SREBF/SREBP | 2.26 | 0.000 |
| REACTOME_Regulation of Cholesterol Biosynthesis by SREBP/SREBF | 2.18 | 0.000 |
| REACTOME_Mitochondrial Translation | 2.16 | 0.000 |
| KEGG_Steroid Biosynthesis | 2.06 | 0.001 |
| KEGG_Spliceosome | 2.04 | 0.001 |
| KEGG_Terpenoid Backbone Biosynthesis | 2.03 | 0.001 |
| REACTOME_Mitochondrial Biogenesis | 2.01 | 0.010 |
| REACTOME_Negative Regulation of MAPK Pathway | 1.96 | 0.018 |
| HALLMARK_TNFa Signaling via NFkB | 1.93 | 0.000 |
| REACTOME_Transcriptional Activation of Mitochondrial Biogenesis | 1.91 | 0.028 |
| HALLMARK_mTORC1 Signaling | 1.86 | 0.001 |
| REACTOME_Respiratory Electron Transport | 1.85 | 0.051 |
| REACTOME_Negative Regulation of NOTCH4 Signaling | 1.81 | 0.056 |
| KEGG_Ribosome | 1.79 | 0.018 |
| REACTOME_Selective Autophagy | 1.79 | 0.062 |
| REACTOME_RAF-Independent MAPK1/3 Activation | 1.78 | 0.061 |
| REACTOME_ATF4 Activates Genes in Response to Endoplasmic Reticulum Stress | 1.78 | 0.060 |
| REACTOME_CTLA4 Inhibitory Signaling | 1.77 | 0.067 |
| REACTOME_RAF Activation | 1.77 | 0.065 |
| REACTOME_Signaling by the B Cell Receptor BCR | 1.76 | 0.066 |
| REACTOME_Butyrate Response Factor 1/BRF1 Binds and Destabilizes mRNA | 1.76 | 0.065 |
| HALLMARK_Myc Targets V1 | 1.75 | 0.003 |
| REACTOME_Regulation of Gene Expression in Beta Cells | 1.74 | 0.071 |
| KEGG_Proteasome | 1.74 | 0.026 |
| KEGG_RNA Degradation | 1.70 | 0.030 |
| KEGG_SNARE Interactions in Vesicular Transport | 1.70 | 0.030 |
| REACTOME_Regulation of RUNX3 Expression and Activity | 1.70 | 0.083 |
| HALLMARK_Hypoxia | 1.69 | 0.005 |
| REACTOME_SHC-Mediated Cascade FGFR4 | 1.69 | 0.085 |
| REACTOME_Energy-Dependent Regulation of mTOR by LKB1 AMPK | 1.68 | 0.090 |
| REACTOME_Downstream Signaling Events of B Cell Receptor/BCR | 1.67 | 0.089 |
| REACTOME_Degradation of Beta Catenin by the Destruction Complex | 1.67 | 0.089 |
| REACTOME_TNFR2 Non-Canonical NF-kB Pathway | 1.67 | 0.090 |
| REACTOME_Signaling by NOTCH4 | 1.67 | 0.088 |
| REACTOME_The Role of GTSE1 in G2/M Progression After G2 Checkpoint | 1.66 | 0.091 |
| KEGG_Long-Term Depression | 1.64 | 0.046 |
| REACTOME_PI3K Cascade FGFR4 | 1.64 | 0.096 |
| REACTOME_TP53 Regulates Metabolic Genes | 1.64 | 0.095 |
| HALLMARK_Androgen Response | 1.63 | 0.008 |
| KEGG_Huntington's Disease | 1.63 | 0.049 |
| KEGG_Cardiac Muscle Contraction | 1.60 | 0.063 |
| REACTOME_Dectin1 Mediated Noncanonical NF-kB Signaling | 1.60 | 0.098 |
| HALLMARK_Oxidative Phosphorylation | 1.58 | 0.010 |
| HALLMARK_Myc Targets V2 | 1.57 | 0.010 |
| KEGG_MAPK Signaling Pathway | 1.57 | 0.073 |
| KEGG_RNA Polymerase | 1.53 | 0.094 |
| HALLMARK_p53 Pathway | 1.48 | 0.025 |
| HALLMARK_PI3k/Akt/mTOR Signaling | 1.33 | 0.080 |
| HALLMARK_Estrogen Response Early | 1.31 | 0.089 |

Table S2. HALLMARK, KEGG and REACTOME Gene Sets Enriched in the iWAT of HF-fed LFABP-/- male mice. FDR < 0.1, |FC| > 1.2

| **Gene Set** | **NES** | **FDR** |
| --- | --- | --- |
| REACTOME_Collagen Degradation | 2.30 | 0.000 |
| REACTOME_Binding and Uptake of Ligands by Scavenger Receptors | 2.30 | 0.000 |
| REACTOME_Assembly of Collagen Fibrils and Other Multimeric Structures | 2.25 | 0.000 |
| REACTOME_Initial Triggering of Complement | 2.21 | 0.000 |
| REACTOME_Collagen Formation | 2.19 | 0.000 |
| REACTOME_Collagen Biosynthesis and Modifying Enzymes | 2.16 | 0.001 |
| REACTOME_Degradation of the Extracellular Matrix | 2.16 | 0.000 |
| REACTOME_Collagen Chain Trimerization | 2.15 | 0.001 |
| KEGG_Renin Angiotensin System | 2.14 | 0.000 |
| REACTOME_Scavenging by Class A Receptors | 2.11 | 0.001 |
| REACTOME_Complement Cascade | 2.09 | 0.001 |
| REACTOME_Activation of Matrix Metalloproteinases | 2.08 | 0.001 |
| REACTOME_MET Activates PTK2 Signaling | 2.07 | 0.002 |
| KEGG_ECM Receptor Interaction | 2.07 | 0.000 |
| REACTOME_Branched-Chain Amino Acid Metabolism | 2.04 | 0.002 |
| REACTOME_Metabolism of Angiotensinogen to Angiotensins | 2.00 | 0.004 |
| REACTOME_MET Promotes Cell Motility | 1.97 | 0.005 |
| KEGG_Lysosome | 1.98 | 0.001 |
| KEGG_Glycosaminoglycan Degradation | 1.97 | 0.001 |
| REACTOME_ECM Proteoglycans | 1.96 | 0.005 |
| KEGG_Hematopoietic Cell Lineage | 1.95 | 0.001 |
| REACTOME_Immunoregulatory Interactions Between a Lymphoid and a Non-Lymphoid Cell | 1.93 | 0.007 |
| REACTOME_Crosslinking of Collagen Fibrils | 1.93 | 0.007 |
| REACTOME_Extracellular Matrix Organization | 1.92 | 0.007 |
| REACTOME_Antimicrobial Peptides | 1.88 | 0.011 |
| REACTOME_Laminin Interactions | 1.86 | 0.014 |
| HALLMARK_Interferon-alpha Response | 1.83 | 0.005 |
| REACTOME_Integrin Cell Surface Interactions | 1.78 | 0.030 |
| REACTOME_Xenobiotics | 1.75 | 0.041 |
| REACTOME_FCGR3A-Mediated IL10 Synthesis | 1.74 | 0.045 |
| REACTOME_Anchoring Fibril Formation | 1.71 | 0.059 |
| REACTOME_Interleukin-10 Signaling | 1.70 | 0.061 |
| KEGG_ABC Transporters | 1.70 | 0.014 |
| KEGG_Propanoate Metabolism | 1.69 | 0.015 |
| REACTOME_O-Linked Glycosylation | 1.68 | 0.073 |
| REACTOME_Glycosphingolipid Metabolism | 1.65 | 0.097 |
| REACTOME_ABC Transporters in Lipid Homeostasis | 1.64 | 0.096 |
| REACTOME_Sialic Acid Metabolism | 1.64 | 0.099 |
| KEGG_Complement and Coagulation Cascades | 1.60 | 0.030 |
| HALLMARK_Epithelial Mesenchymal Transition | 1.57 | 0.063 |
| KEGG_Sphingolipid Metabolism | 1.53 | 0.048 |
| KEGG_Valine, Leucine and Isoleucine Degradation | 1.52 | 0.050 |
| KEGG_Other Glycan Degradation | 1.47 | 0.064 |
| HALLMARK_Coagulation | 1.47 | 0.080 |
| KEGG_Metabolism of Xenobiotics by Cytochrome P450 | 1.43 | 0.085 |
| KEGG_O-Glycan Biosynthesis | 1.41 | 0.098 |
| HALLMARK_Xenobiotic Metabolism | 1.39 | 0.099 |

Table S3. HALLMARK, KEGG and REACTOME Gene Sets Downregulated in the iWAT of HF-fed LFABP-/- male mice. FDR < 0.1, |FC| > 1.2

1.	A. M. Gajda *et al.*, Direct Comparison of Mice Null for Liver or Intestinal Fatty Acid-binding Proteins Reveals Highly Divergent Phenotypic Responses to High Fat Feeding. **288**, 30330-30344 (2013).
